# Neural responses to natural and model-matched stimuli reveal distinct computations in primary and non-primary auditory cortex

**DOI:** 10.1101/419168

**Authors:** Sam V. Norman-Haignere, Josh H. McDermott

## Abstract

A central goal of sensory neuroscience is to construct models that can explain neural responses to complex, natural stimuli. As a consequence, sensory models are often tested by comparing neural responses to natural stimuli with model responses to those stimuli. One challenge is that distinct model features are often correlated across natural stimuli, and thus model features can predict neural responses even if they do not in fact drive them. Here we propose a simple alternative for testing a sensory model: we synthesize stimuli that yield the same model response as a natural stimulus, and test whether the natural and “model-matched” stimulus elicit the same neural response. We used this approach to test whether a common model of auditory cortex – in which spectrogram-like peripheral input is processed by linear spectrotemporal filters – can explain fMRI responses in humans to natural sounds. Prior studies have that shown that this model has good predictive power throughout auditory cortex, but this finding could reflect stimulus-driven correlations. We observed that fMRI voxel responses to natural and model-matched stimuli were nearly equivalent in primary auditory cortex, but that non-primary regions showed highly divergent responses to the two sound sets, suggesting that neurons in non-primary regions extract higher-order properties not made explicit by traditional models. This dissociation between primary and non-primary regions was not clear from model predictions due to the influence of stimulus-driven response correlations. Our methodology enables stronger tests of sensory models and could be broadly applied in other domains.

**Author Summary:** Modeling neural responses to natural stimuli is a core goal of sensory neuroscience. Here we propose a new approach for testing sensory models: we synthesize a “model-matched” stimulus that yields the same model response as a natural stimulus, and test whether it produces the same neural response. We used model-matching to test whether a standard model of auditory cortex can explain human cortical responses measured with fMRI. Model-matched stimuli produced nearly equivalent voxel responses in primary auditory cortex, but highly divergent responses in non-primary regions. This dissociation was not evident using more standard approaches for model testing, and suggests that non-primary regions compute higher-order stimulus properties not captured by traditional models. The methodology could be broadly applied in other domains.

## Introduction

One definition of understanding a neural system is to be able to build a model that can predict its responses. Responses to natural stimuli are of particular interest, both because natural stimuli are complex and varied, and thus provide a strong test of a model, and because sensory systems are presumably adapted to represent features present in natural stimuli^1–3^. The evaluation of models by their ability to predict responses to natural stimuli is now widespread in sensory neuroscience^4–15^.

A challenge for this approach is that because natural stimuli are richly structured, the features of a set of natural stimuli in one model (or model stage) are often correlated with the features in other models (or model stages)^16^. Model features can thus in principle predict neural responses to a natural stimulus set even if the neural responses are in fact driven by other features not captured by the model. Related issues have been widely discussed in the receptive field estimation literature^4,17^, but have been less noted in cognitive neuroscience^16,18^.

A canonical example of this phenomenon occurs in the auditory domain, where there is still considerable uncertainty regarding computational descriptions of cortical processing. Consider a common model of auditory processing, in which a sound waveform is processed by two stages of filters intended to mimic cochlear and cortical filtering, respectively^19^ (Figure 1A). The filters in the second model stage are tuned to temporal and spectral modulations in the spectrogram-like representation produced by the cochlea. Such filters and variants thereof are commonly used to account for human perceptual abilities^20–23^ and to explain neural responses throughout the auditory pathway^2,7,9,11,12,24–35^. But in natural stimuli, the responses of these second-stage filters are often correlated with other sound properties, such as semantic categories (Figure 1B)^36^, which can confound the interpretation of neural responses. Speech, for instance, has a distinctive temporal modulation rate that corresponds loosely to the rate of syllabic patterning^37^, music has distinctive temporal modulations reflective of its beat structure^38^, and both speech and music have characteristic spectral modulations due to harmonic frequency structure^19^. However, speech, music, and other natural sounds also have many unique properties that are not captured by spectrotemporal modulation alone^39^. Thus, if a neuron responds more to speech than to other sounds, modulation filters may be able to predict the neuron’s response, even if the response is driven by another property of speech that is not captured by such filters. This is what we term a “stimulus-driven response correlation”, created when different stimulus properties (e.g. spectrotemporal modulations and semantic categories) are correlated within a particular stimulus set, making their contribution to the neural response difficult to tease apart.

**Figure 1.**
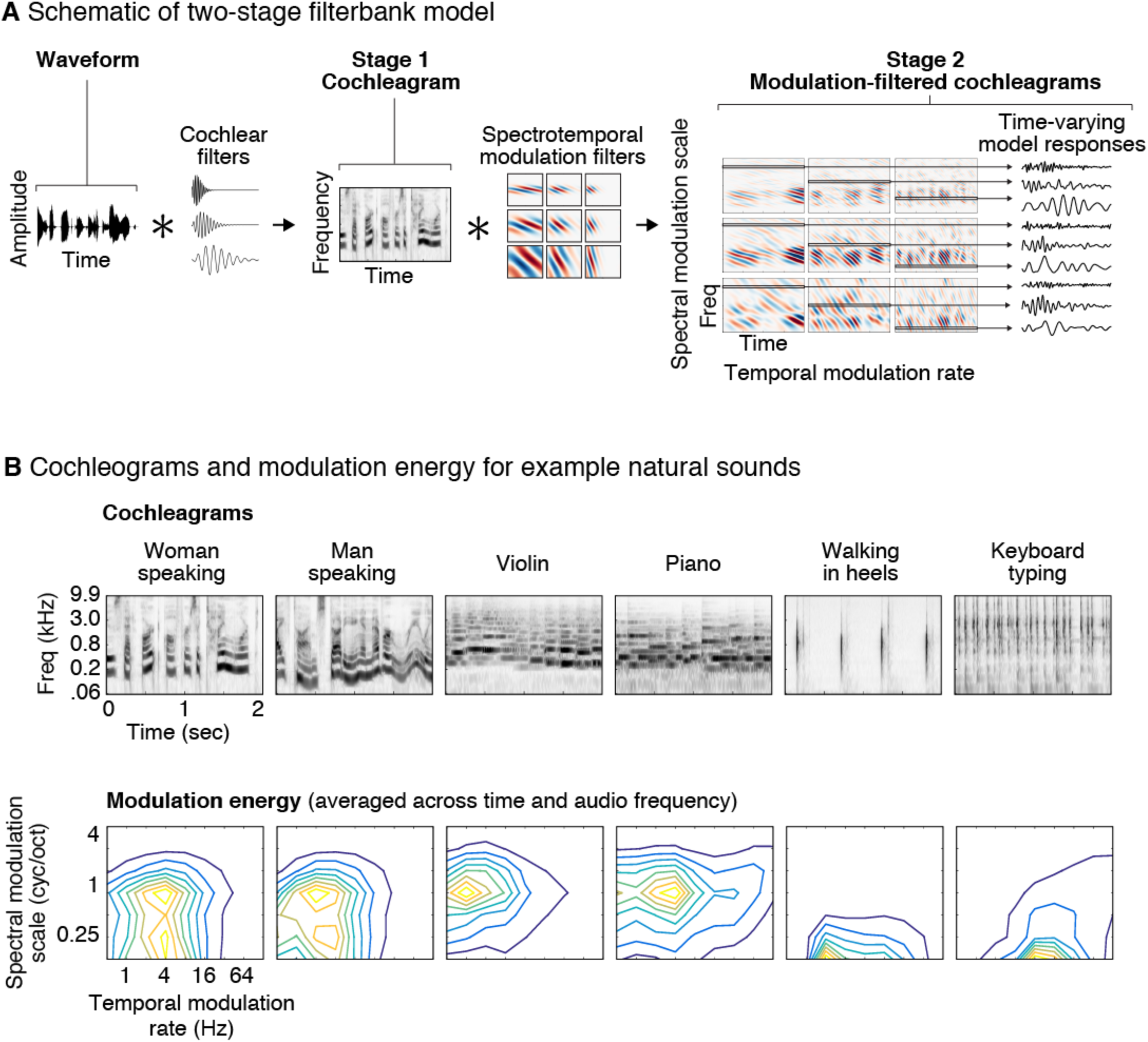
Illustration of the auditory model tested in this study. (A) The model consists of two cascaded stages of filtering. In the first stage, a cochleagram is computed by convolving each sound with audio filters tuned to different frequencies, extracting the temporal envelope of the resulting filter responses, and applying a compressive nonlinearity to simulate the effect of cochlear amplification (for simplicity, envelope extraction and compression are not illustrated in the figure). The result is a spectrogram-like output that represents sound energy as a function of time and frequency. In the second stage, the cochleagram is convolved in time and frequency with filters that are tuned to different rates of temporal and spectral modulation. The output of the second stage can be conceptualized as a set of filtered cochleagrams, each highlighting modulations at a particular temporal rate and spectral scale. Each frequency channel of these filtered cochleagrams represents the time-varying output of a single model feature that is tuned to audio frequency, temporal modulation rate, and spectral modulation scale. (B) Cochleagrams and modulation spectra are shown for six example natural sounds. Modulation spectra plot the energy (variance) of the second-stage filter responses as a function of temporal modulation rate and spectral modulation scale, averaged across time and audio frequency. Different classes of sounds have characteristic modulation spectra.

Here we propose a complementary method for evaluating models that circumvents the challenge of stimulus-driven response correlations. The idea is simple: we synthesize stimuli that yield the same response in a model as a natural stimulus, and then test whether the “model-matched” stimulus elicits the same neural response as the natural stimulus. The synthesized sounds are not influenced by the correlations between different feature sets that may exist in natural stimuli because they are constrained only by the features in the model. As a result, they generally differ in other properties that could potentially be important to the neural response, and often sound markedly different from their natural counterparts. Comparing responses to natural and model-matched sounds thus provides a strong test of the model’s explanatory power.

We demonstrate the method by using it to evaluate whether a common filter bank model of the auditory cortex can explain human cortical responses to natural sounds measured with fMRI. Many prior fMRI studies of auditory cortex have identified aspects of cortical tuning that are unique to non-primary regions^16,40,41^, such as selectivity for voice^42^, speech^43,44^ and music^45–47^. At the same time, other studies have demonstrated that the standard filter bank model has good predictive accuracy throughout primary and non-primary regions^7,12,41,45^, raising the possibility that primary and non-primary regions encode sound using similar representations. Alternatively, such predictions could in part reflect stimulus-driven correlations. Here, we addressed this question by comparing cortical fMRI responses to natural and model-matched stimuli.

The model-matched stimuli were synthesized to yield the same response as a natural sound in one of several models of varying complexity, ranging from a model of just the cochlea’s response to the two stage spectrotemporal filter-bank model shown in Figure 1A^19^. Our results show that tuning for temporal and spectral modulations explains much of the voxel response to natural sounds in human primary auditory cortex (PAC), but much less of the response in non-primary areas. This functional difference between primary and non-primary regions was much less evident using conventional model predictions due to the effect of stimulus-driven response correlations. Our findings provide novel evidence for functional differentiation between primary and non-primary auditory cortex, and suggest that non-primary regions build higher-order representations that cannot be explained by standard models. Our methodology could provide stronger tests of neural models in any system for which models are used to predict neural responses.

## Results

### Overview of model-matching method and underlying assumptions

The goal of this paper was to test whether conventional auditory models can explain voxel responses in auditory cortex to natural sounds. The models we consider are described by a set of model features (*m* _*k*_ (*t*)) each of which has a time-varying response to sound determined by the feature’s filter (Figure 1A&2A). In general, the response of these features will differ across natural sounds, both in their temporal pattern and their time-averaged properties (Figure S1A). The BOLD signal reflects a time-averaged measure of neural activity, and thus we expect that any two sounds with the same time-averaged properties (i.e. the same histogram of amplitudes) should yield the same fMRI response even if the temporal pattern is different. To test this prediction, we iteratively modified a noise stimulus, which was initially unstructured, so as to match the histogram of each filter’s response (Figure S1B), similar to methods for texture synthesis^39,48–50^. Because the temporal patterns of the model responses are unconstrained, the model-matched sounds differ from the natural sounds they were matched to.

Formally, we assume that the response of a voxel to a sound can be approximated as the weighted sum of time-averaged neuronal firing rates. Here, we assume the voxel response to be a single number because the sounds we present are short relative to the time scale of the BOLD response. Our goal is to test whether these model feature responses approximate neuronal responses within a voxel, in which case we should be able to approximate the voxel’s response (v_*i*_) as a weighted sum of time-averaged model responses (a_*k*_) (Figure 2A):

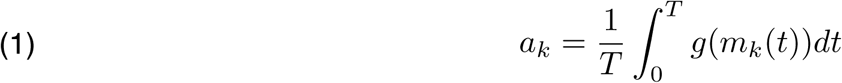

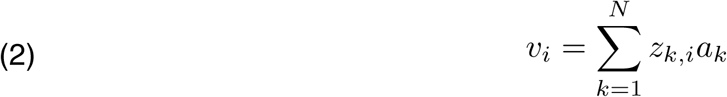

where *g* is an (unknown) point-wise function that maps the model responses to a neuronal firing rate (e.g. a rectifying nonlinearity), *z*_*k,i*_ is the weight of model feature *k* in voxel *i*, and T is the duration of the response to a sound. The most common approach for testing equations 1 and 2 is to estimate the weights (*z*_*k,i*_,) that best predict a given voxel’s response to natural sounds (for a particular choice of *g*), and to assess the cross-validated prediction accuracy of the model using these weights (via explained variance). Here we instead test the above equations by synthesizing a ‘model-matched’ sound that should yield the same voxel response as a natural sound for all voxels that are accurately described by the model (Figure 2A). We then test the model’s validity by assessing whether the voxel responses to the two sounds are similar.

**Figure 2.**
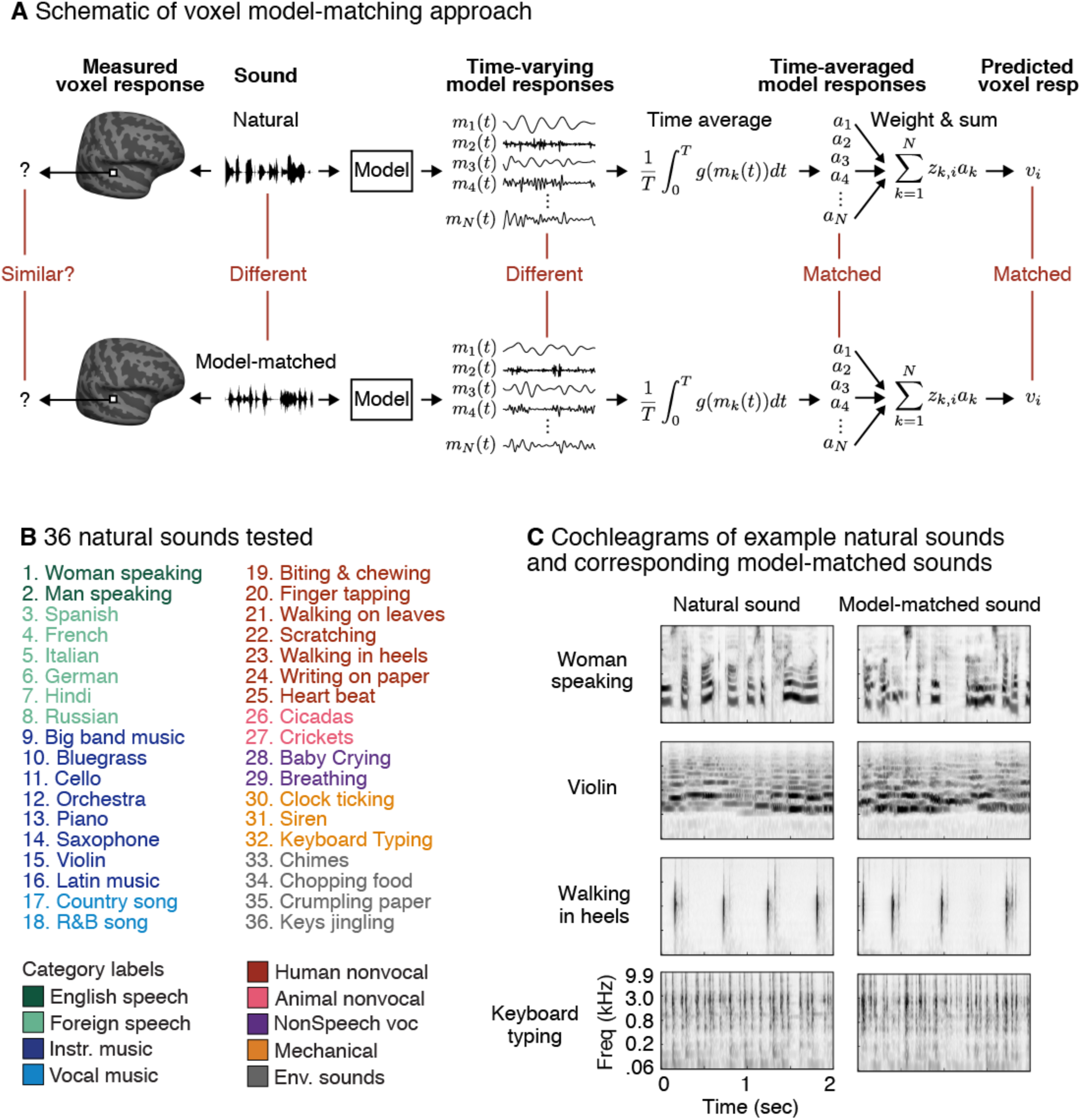
Model-matching methodology and experimental stimuli. (A) The logic of the model-matching procedure as applied to fMRI. The models we consider are defined by the time-varying response of a set of model features (*m*_*k*_(*t*)) to a sound (as in the auditory model shown in Figure 1A). Because fMRI is thought to pool activity across neurons and time, we modeled fMRI voxel responses as weighted sums of time-averaged model responses (equations 1-2) (*a*_*k*_ corresponding to the time-averaged model responses and *z*_*k,i*_ to the weight of model feature *k* in voxel *i*). Model-matched sounds were designed to produce the same time-averaged response for all of the features in the model (all *a*_*k*_ matched), and thus to yield the same voxel response regardless of how these time-averaged activities are weighted (for voxels containing neurons that can be approximated by the model features). The temporal response pattern of the model features was otherwise unconstrained. As a consequence the model-matched sounds were distinct from the natural sounds they were matched to. (B) Stimuli were derived from a set of 36 natural sounds. The sounds were selected to produce high response variance in auditory cortical voxels, based on the results of a prior study^45^. Font color denotes membership in one of 9 semantic categories (as determined by human listeners^45^. (C) Cochleagrams are shown for four natural and model-matched sounds constrained by the spectrotemporal modulation model shown in Figure 1A^18^.

In principle, one could synthesize a separate model-matched sound for each voxel after learning the weights (*z*_*k,i*_). However, this approach is impractical given the many thousands of voxels in auditory cortex. Instead, we matched the time-averaged response of all features in the model (i.e. all *a*_*k*_ in equation 2 are matched; see Figure 2A), which guarantees that all voxel responses should be matched regardless of that voxel’s weights. We accomplished this objective by matching the histogram of each feature’s response (Figure S1; see *Model-matching synthesis algorithm* in the Methods)^48^. Histogram matching implicitly equates the time-averaged response of the model features for any point-wise transformation (*g*) (since for any such transformation, the time-averaged response can be approximated via its histogram), and thus obviates the need to choose a particular nonlinearity.

Whether or not a voxel responds similarly to natural and model-matched sounds depends on the response properties of the model features and underlying neurons. If the model features are good approximations to the neurons in a voxel, then the voxel response to natural and model-matched sounds should be similar; if not, they could differ. Here we consider model features that are tuned to different patterns of temporal and/or spectral modulation^19^ in a spectrogram-like representation of sound termed a cochleagram (Figure 1A), produced by passing a sound signal through filters designed to mimic cochlear tuning. Each model feature is associated with a time-frequency filter tuned to a particular temporal rate and/or scale, as well as to a particular audio frequency. The response of each model feature is computed by convolving the spectrotemporal filter with the cochleagram.

Although the response timecourses of the models considered here are sufficient to reconstruct the stimulus with high accuracy, the time-averaged properties of the filters, as captured by a histogram, are not. As a consequence the model-matched sounds differed from the natural sounds they were matched to. Indeed many of the model-matched stimuli sound unnatural (see http://mcdermottlab.mit.edu/svnh/model-matching/Stimuli_from_Model-Matching_Experiment.html for examples). This observation demonstrates that the time-averaged properties of the model’s features, which approximately capture the modulation spectrum (Figure 2A), fail to capture many perceptually salient properties of natural stimuli (e.g. the presence of phonemic structure in speech or melodic contours in music). This additional structure is conveyed by temporal patterns in the feature responses, which are not made explicit by the model, but which might be extracted by additional layers of processing not present in modulation filter models. If the neurons in a voxel respond to such higher-order properties (e.g. the presence of a phoneme or melodic contour), we might expect their time-averaged response to differ between natural and model-matched sounds. Thus, by measuring the similarity of voxel responses to natural and model-matched sounds, we can test whether the features of the filter bank model are sufficient to explain their response, or whether other features are needed.

### Comparing fMRI responses to natural and model-matched sounds

We measured fMRI responses to a diverse set of 36 natural sounds and their corresponding model-matched sounds (Figure 2B) (Chi et al. 2005). Each sound was originally 10 seconds in duration, but the sounds were broken up into successively presented 2-second excerpts to accommodate the fMRI scanning procedure (Figure S2; see *Stimulus presentation and scanning procedure* in the Methods). The model-matched sounds were constrained by all of the features from the two-stage filter bank model shown in Figure 1A (later we show results from sounds constrained by simpler models; see *Model representation* in the Methods). We first plot the response of two example voxels from a single subject (Figure 3A), which illustrate some of the dominant trends in the data. One voxel was located in the low-frequency area of the “high-low-high” tonotopic gradient thought to span PAC, and which is organized in a roughly V-shaped pattern^51–55^. Another voxel was located outside of tonotopically defined PAC. We note that how best to define PAC is a matter of activate debate^54,56–58^, and thus we have quantified our results using both tonotopic and anatomical definitions of PAC (described below).

**Figure 3.**
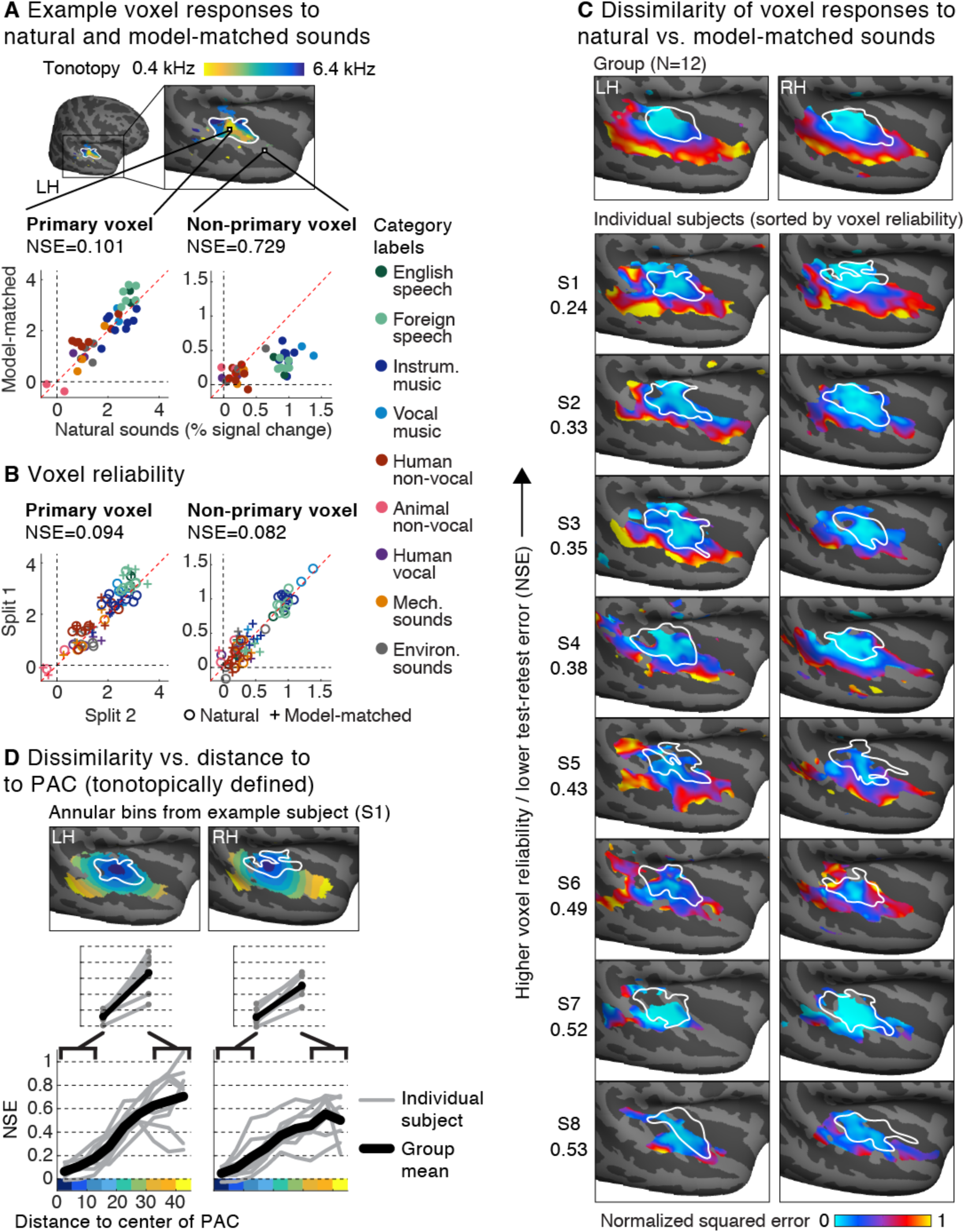
Voxel responses to natural and model-matched sounds. (A) Responses to natural and model-matched sounds from two example voxels from a single subject. One voxel is drawn from the low-frequency region of primary auditory cortex (defined tonotopically) and one from outside of PAC. A tonotopic map measured in the same subject is shown for anatomical comparison; the map plots the pure tone frequency that produced the highest voxel response. Each dot represents the response to a single pair of natural and model-matched sounds. The primary voxel responded similarly to natural and model-matched sounds, while the non-primary voxel exhibited a weaker response to model-matched sounds. We quantified the dissimilarity of voxel responses to natural and model-matched sounds using a normalized squared error metric (NSE) (see text for details). (B) Split-half reliability of the responses to natural (circles) and model-matched sounds (crosses) for the two voxels shown in panel A. Both primary and non-primary voxels exhibited a reliable response (and thus a low NSE between the two measurements). (C) Maps plotting the NSE between each voxel’s response to natural and model-matched sounds, corrected for noise in fMRI measurements (see Figure S4 for uncorrected maps). Maps are shown both for voxel responses from eight individual subjects (who were scanned more than the other subjects) and for group responses averaged across 12 subjects in standardized anatomical coordinates (top). The white outline plots the boundaries of primary auditory cortex, defined tonotopically. Only voxels with a reliable response are plotted (see text for details). Subjects are sorted by the median test-retest reliability of their voxel responses in auditory cortex, as measured by the NSE. (D) A summary figure plotting the dissimilarity of voxel responses to natural and model-matched sounds as a function of distance to the low-frequency region of PAC (see Figure S5 for an anatomically-based analysis). This figure was computed from the individual subject maps shown in panel C. Voxels were binned based on their distance to PAC in 5 mm intervals. The bins for one example subject (S1) are plotted. Each gray line represents a single subject (for each bin the median NSE value across voxels is plotted), and the black line represents the average across subjects. Primary and non-primary auditory cortex were defined as the average NSE value across the three bins closest and furthest from PAC (inset). In every subject and hemisphere, we observed larger NSE values in non-primary regions.

As shown in Figure 3A, the response of the primary voxel to natural and model-matched sounds was similar. By contrast, the non-primary voxel responded notably less to the model-matched sounds. We quantified the dissimilarity of responses to natural and model-matched sounds by computing the squared error between corresponding pairs of natural and model-matched sounds, normalized by the squared error that would be expected if there was no correspondence between the two sound sets (see *Normalized squared error* in the Methods). We quantified response differences using the squared error rather than the correlation because model-matching makes no prediction for how responses to natural and model-matched sounds should differ if the model is inaccurate, and in practice responses to model-matched sounds were often weaker in non-primary regions, a phenomena that would not have been captured by correlation. At the end of the results, we quantify how natural and model-matched sounds differ.

We found that the NSE was higher for the non-primary voxel (NSE=0.729) than the primary voxel (NSE=0.101), reflecting the fact that the non-primary voxel showed a more dissimilar response to natural and model-matched sounds. Moreover, most of the error between responses to natural and model-matched sounds in the primary voxel could be attributed to noise in the fMRI measurements, since a similar NSE value was observed between two different measurements of the voxel’s response to natural sounds (NSE=0.094) (Figure 3B). By contrast in the non-primary voxel, the test-retest NSE (NSE=0.082) was much lower than the NSE between responses to natural and model-matched sounds, indicating that the difference in response to natural and model-matched sounds cannot be explained by lower SNR.

We quantified these effects across voxels by plotting the NSE between responses to natural and model-matched sounds for each voxel (Figures 3C). Maps were computed from voxel responses in eight individual subjects who were scanned substantially more than the other subjects (see *Participants* in the Methods for details) and from responses that were averaged across all twelve subjects after aligning their brains. Data were collected using two different experiment paradigms that differed in the sounds that were repeated within a scanning session. The results were similar between the two paradigms (Figure S3), and so we describe them together (see Methods for details; subjects S1, S2, S3, S7, S8 were scanned in Paradigm I; subjects S4, S5, S6 were scanned in Paradigm II; group results are based on data from Paradigm I; S1 participated in both paradigms but only data from Paradigm II is shown). In Paradigm I, only responses to natural sounds were repeated, while in Paradigm II both natural and model-matched sounds were repeated. Only voxels with a reliable response are plotted (test-retest NSE < 0.4; see *Evaluating the noise-corrected NSE with simulated data* in the Methods for a justification of this criterion; reliability was calculated using natural sounds for Paradigm I and both natural and model-matched sounds for Paradigm II). Subjects have been ordered by the overall reliability of their data (median test-retest NSE across the superior temporal plane and gyrus, evaluated using natural sounds so that we could apply the same metric to subjects from Paradigms I and II). These maps have been corrected for noise in the fMRI measurements (see *Noise-correcting the normalized squared error* in the Methods), but the results were similar without correction (Figure S4).

Both group and individual subject maps revealed a substantial change across the cortex in the similarity of responses to natural and model-matched sounds. Voxels in PAC showed a similar response to natural and model-matched sounds with noise-corrected NSEs approaching 0, indicating nearly identical responses. Moving away from PAC, NSE values rose substantially, reaching values near 1 in regions far from PAC. This pattern of results suggests that the filter bank model can explain much of the voxel response in primary regions, but much less of the response in non-primary regions, plausibly because non-primary regions respond to higher-order features not made explicit by the model. This result is suggestive of a hierarchy of feature selectivity in auditory cortex, and demonstrates where in the cortex the standard filter bank model fails to explain voxel responses.

We quantified the gradient we observed between primary and non-primary voxels by binning the NSE of voxels from individual subjects based on their distance to PAC. Similar results were observed for tonotopic (Figure 3D) and anatomical definitions of PAC (Figure S5; PAC was defined either as the center of the high-low-high gradient, or as the center of anatomical region TE1.1^58^ which is located in posteromedial HG). To directly compare primary and non-primary regions we then averaged NSE values within the three bins nearest and furthest from PAC (Figure 3D, inset). This analysis revealed that responses to natural and model-matched sounds became more dissimilar in non-primary regions in both the left and right hemisphere of every subject tested, leading to a highly significant difference between primary and non-primary regions (t(7) > 6.79, p < 0.001 for both hemispheres and for both tonotopic and anatomical definitions of PAC). The gradient between primary and non-primary regions was observed in both scanning paradigms regardless of smoothing (Figure S3), and could not be explained by selectivity for intelligible speech (a similar pattern was observed when intelligible speech sounds were excluded from the analysis; see Figure S6). These results also could not be explained by variations in voxel reliability across brain regions: both because our NSE measures were noise-corrected, and because voxel responses were similarly reliable throughout primary and non-primary regions (Figure S4C). As a consequence the increase in the NSE between natural and model-matched sounds between primary and non-primary regions was significantly greater than the change in voxel reliability. This was true using both corrected and uncorrected values for the natural vs. model-matched NSE, both tonotopic and anatomical definitions of PAC, and with reliability measured using just natural sounds for Paradigm I and both natural and model-matched sounds for Paradigm II (t(7) > 6.41; p < 0.001; see Figure S3 for a breakdown by paradigm). Thus, our results demonstrate that the modulation filter bank model is worse at accounting for voxel responses in non-primary regions.

### Comparing responses to sounds matched on subsets of model features

We next used a similar approach to test whether responses in PAC could be explained by simpler models. For example, if neurons in a voxel are tuned primarily to audio frequency, than any sound with a similar spectrum should produce a similar response, regardless of its modulation properties. To test such alternative models, we synthesized three new sounds for each natural sound. Each synthetic sound was matched on a different subset of features from the full model (Figure 4A). One sound was synthesized to have the same marginal distribution of cochlear envelopes as a natural sound, and thus a similar audio spectrum, but its modulation properties were otherwise unconstrained. Another sound was constrained to have the same temporal modulation statistics within each cochlear frequency channel, computed using a bank of modulation filters modulated in time but not frequency. A third sound was synthesized to have matched spectral modulation statistics, computed from a bank of filters modulated in frequency but not time. All of the modulation-matched sounds also had matched cochlear marginal statistics, thus making it possible to test whether adding modulation structure enhanced the similarity of cortical responses to natural and model-matched sounds.

**Figure 4.**
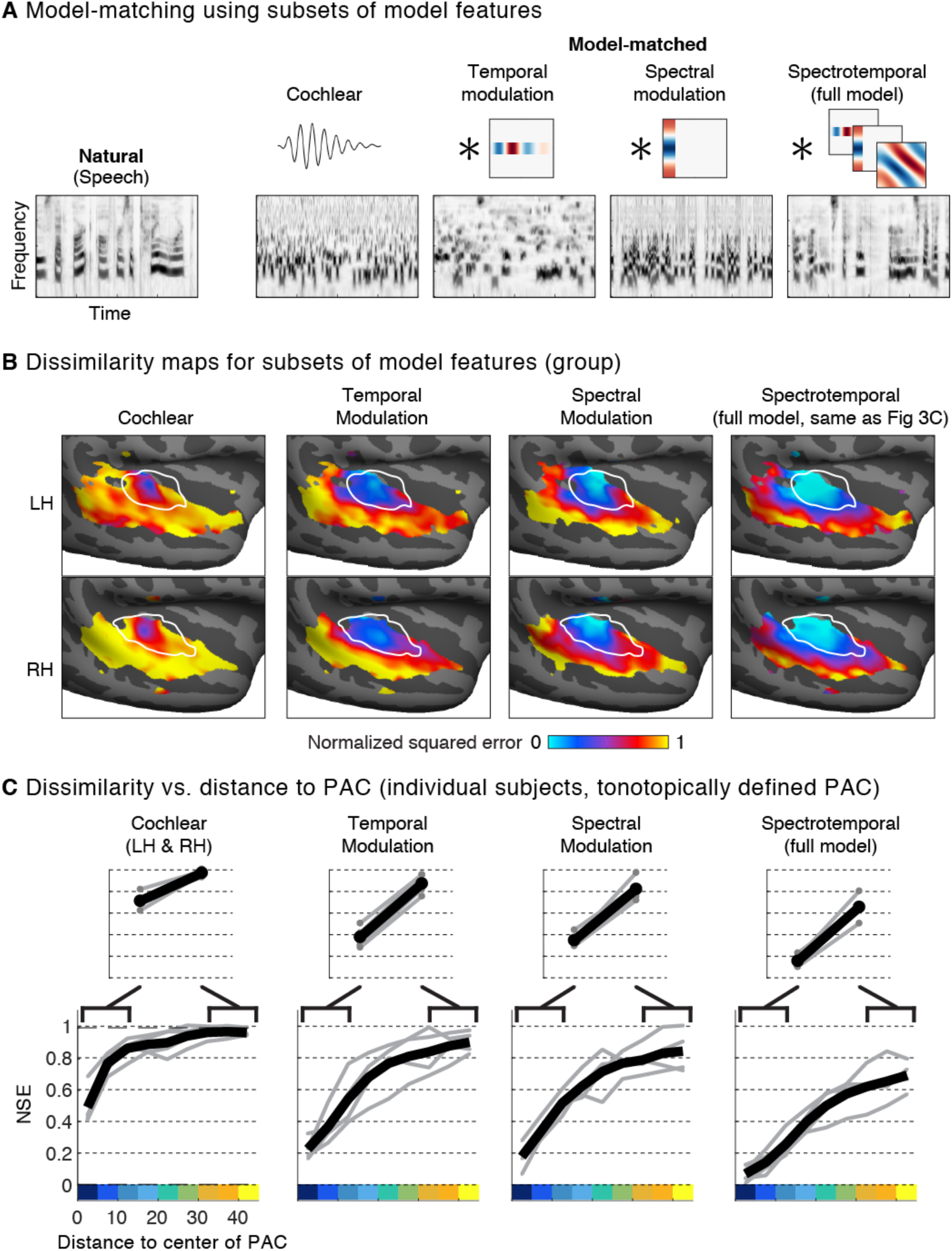
Comparison of responses to model-matched sounds constrained by different models. (A) Cochleagrams for an example natural sound and several corresponding model-matched sounds constrained by subsets of features from the full two-stage model. Cochlear-matched sounds were constrained by time-averaged statistics of the cochleagram representation, but not by any responses from the second stage filters. As a consequence, they had a similar spectrum and overall depth of modulation as the corresponding natural sound, but were otherwise unconstrained. The other three sounds were additionally constrained by the response of second-stage filters, tuned either to temporal modulation, spectral modulation, or both temporal and spectral modulation (the full model used in Figure 3). Temporal modulation filters were convolved separately in time with each cochlear frequency channel. Spectral modulation filters were convolved in frequency with each time-slice of the cochleagram. The absence of spectral modulation filters causes the frequency channels to become less correlated, while the absence of temporal modulation filters results in a signal with more rapid temporal variations than that present in natural speech. (B) Maps of the normalized squared error between responses to natural and model-matched sounds, constrained by each of the four models. Format is the same as panel 3C. See Figure S7 for maps from individual subjects. (C) Dissimilarity between responses to natural and model-matched vs. distance to the low-frequency area of PAC. Format is the same as panel 3D. Results are based on data from the four subjects that participated in Paradigm I, because model-matched sounds constrained by subsets of features were not tested in Paradigm II.

The results of this analysis suggest that all of the model features are necessary to account for voxel responses to natural sounds in primary auditory cortex (Figure 4B&C; Figure S7). Responses to model-matched sounds constrained just by cochlear statistics differed substantially from responses to natural sounds even in PAC, leading to significantly larger NSE values than those observed for the full model (p < 0.001 via bootstrapping across subjects; see *Statistics* in the Methods). Thus, even though PAC exhibits selectivity for frequency due to tonotopy, this selectivity only accounts for a small fraction of its response to natural sounds. Responses to natural and model-matched sounds in PAC became more similar when the sounds were constrained by either temporal or spectral modulation properties alone (NSE temporal < NSE cochlear: p < 0.001 via bootstrapping). However, we only observed NSE values near 0 when sounds were matched in both their temporal and spectral modulation properties (NSE full model < NSE temporal: p < 0.001; NSE full model < NSE spectral: p < 0.001). These results provide further support for the idea that selectivity for both temporal and spectral modulation is a prominent feature of cortical tuning in PAC^7,30,31^.

### Predicting responses to natural sounds from model features

Part of the motivation for using model-matched stimuli comes from the more common approach of predicting responses to natural stimuli from the features of a model (e.g. via linear regression). As discussed above, good predictive accuracy is not sufficient to guarantee that the features of a model drive a neural response, due to the potential for correlations between different feature sets across natural stimuli. Model-matching provides one way to circumvent this issue, since the synthesized sounds are only constrained by the statistics of the particular model being tested. Here, we test whether our approach yields novel insights compared with simply predicting cortical responses to natural sounds from model features.

We attempted to predict responses to the 36 natural sounds from time-averaged statistics of the same model features used to generate the model-matched sounds (Figure 5A; see Figure S8 for individual-subject prediction error maps for the full spectrotemporal model). Specifically, we used ridge regression to predict voxel responses from the amplitude of each model feature’s response to each natural sound^6,7^, measured as the standard deviation across time (for cochlear features, we used the mean rather than the standard deviation because the features had already been passed through an envelope extraction operation, and the mean thus conveys the amplitude of the filter’s response). Histogram matching implicitly matches all time-averaged statistics of a distribution, including the standard deviation. Predictions based on a single time-averaged statistic, like the standard deviation, therefore provide a conservative estimate of the predictive power of time-averaged statistics. Good predictions in voxels whose responses to model-matched sounds deviated from those to natural sounds would thus suggest that prediction-based analyses overestimate of the model’s explanatory power. We quantified prediction accuracy by measuring the NSE between measured and predicted responses for left-out sounds that were not used to learn the regression weights (see *Model predictions* in Methods).

**Figure 5.**
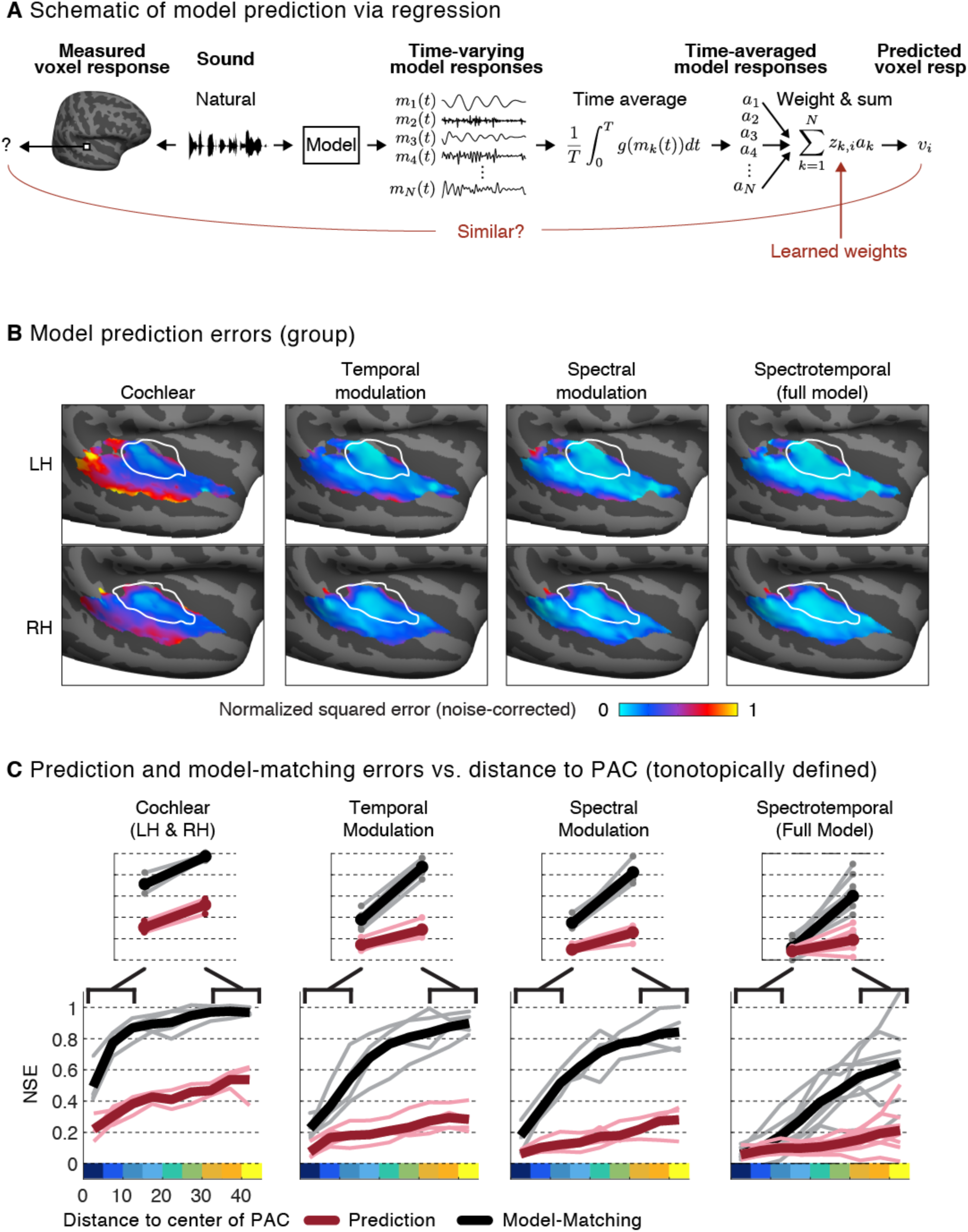
Predicted responses to natural sounds via regression using the same auditory model as that used to constrain the model-matched sounds. (A) Schematic of regression procedure used to predict neural responses from model features. For each natural sound, we computed the response timecourse for each feature in the model, as was done for model-matching. We then computed a time-averaged measure of each feature’s activity (the mean across time for the cochlear features, because they are the result of an envelope operation, and the standard deviation for the modulation features, because they are raw filter outputs), and estimated the weighted combination of these time-averaged statistics that yielded the best-predicted response (using ridge regression, cross-validated across sounds). (B) Maps showing the prediction error (using the same NSE metric employed in Figures 3&4) between measured and predicted responses to natural sounds for the corresponding models shown in Figure 4 (see Figure S8 for maps from individual subjects). (C) Prediction error vs. distance to the low-frequency area of PAC (maroon lines, thin lines correspond to individual subjects, thick lines correspond to the group average). For comparison, the corresponding NSE values derived from the model-matching procedure are re-plotted from Figure 4C (black lines). The analyses are based on individual subject maps. Results for the full-model (rightmost plot) are based on data from the same 8 subjects shown in Figure 3C. Results for model subsets (cochlear, temporal modulation and spectral modulation) are based on data from 4 subjects that were scanned in Paradigm I (sounds constrained by subsets of model features were not tested in Paradigm II).

Overall, we found that that voxel responses to natural sounds were substantially more similar to the predicted model responses than to the measured responses to the model-matched stimuli (Figure 5B&C), leading to smaller NSEs for model predictions compared with model-matching. This difference was particularly pronounced in non-primary regions, where we observed relatively good predictions from the full two-stage model despite highly divergent responses to model-matched sounds, leading to a significant interaction between the type of model evaluation (model prediction vs. model matching) and region (primary vs. non-primary) (t(7)>8.02, p < 0.001 for both tonotopic and anatomical definitions of PAC). Because the natural and model-matched sounds were matched in the features used for prediction, the divergent responses to the two sound sets implies that the features used for prediction do not in fact drive the response. Thus, good predictions for natural sounds in the presence of divergent model-matched responses must reflect the indirect influence of correlations between the features of the model and the features that actually drive the neuronal response. Model-matching thus reveals a novel aspect functional organization not clearly evident from model predictions, by demonstrating the failure of the filter bank model to account for non-primary responses.

Our prediction analyses were based on responses to 36 natural sounds which is small relative to the sound sets that have been used elsewhere to evaluate model predictions^7,45,59^. Because our analyses were cross-validated, small sound sets should reduce prediction accuracy, and thus cannot explain our finding that model predictions were better than would be expected given responses to model-matched sounds. Nonetheless, we assessed the robustness of our findings by also predicting responses to a larger set of 165 natural sounds^45^. We observed similar results with this larger sound set, with relatively good prediction accuracy for the full spectrotemporal model throughout primary and non-primary auditory cortex (Figure S9).

Another way to assess the utility of the model-matching approach is to train a model to predict natural sounds, and then test its predictive accuracy on model-matched sounds (and vice-versa). In practice, this approach yielded similar results to directly comparing responses to natural and model-matched sounds: good cross-predictions in primary auditory cortex, but poor cross-predictions in non-primary auditory cortex (Figure S10). This observation is expected given that a) the model predictions for natural sounds were good throughout auditory cortex and b) responses to natural and model-matched sounds diverged in non-primary regions, but provides a consistency check of the two types of analyses.

### Voxel decomposition of responses to natural and model-matched sounds

All of our analyses thus far have been performed on individual voxels, summarized with maps plotting the normalized squared error between each voxel’s response to natural and model-matched sounds. But these error maps do not reveal in what respect the responses to natural and model-matched sounds differ, and due to the large number of voxels it is not feasible to simply plot all of their responses. We previously found that voxel responses to natural sounds can be approximated as a weighted sum of a small number of canonical response patterns (components)^45^ (Figure 6A). Specifically, six components explained over 80% of the noise-corrected response variance to a diverse set of 165 natural sounds across thousands of voxels. We thus used these six components to summarize the responses to natural and model-matched sounds described here. This analysis was possible because many of the subjects from this experiment also participated in our prior study. As a consequence, we were able to learn a set of voxel weights that reconstructed the component response patterns from our prior study, and then apply these same weights to the voxel responses from this experiment (see *Voxel decomposition* in the Methods).

**Figure 6.**
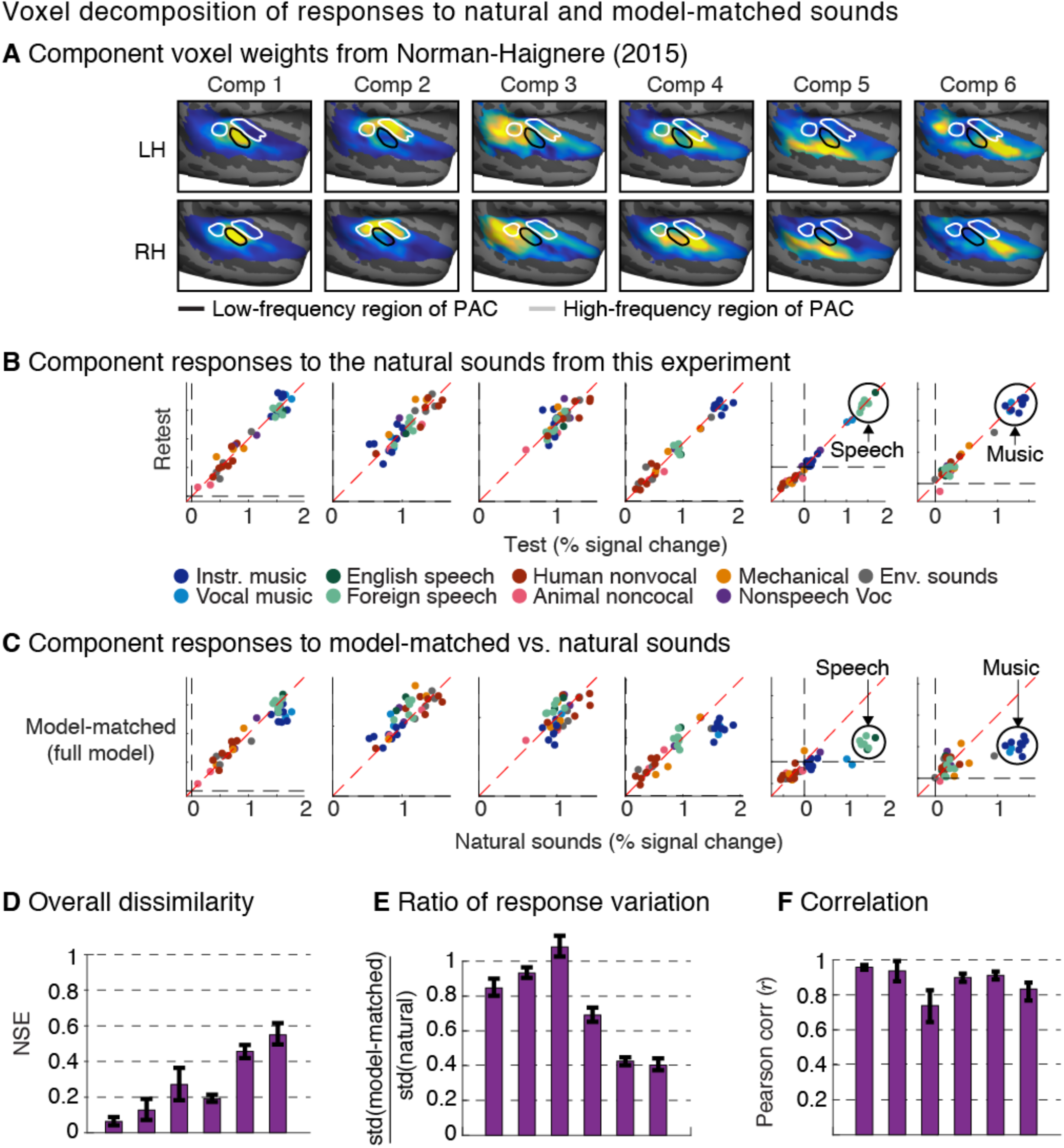
Voxel decomposition of responses to natural and model-matched sounds. Previously we found that much of the voxel response variance to natural sounds can be approximated as a weighted sum of six canonical response patterns (“components”)^45^. This figure shows the response of these components to the natural and model-matched sounds from this experiment. (A) The group components weights from Norman-Haignere et al. (2015)^45^ are re-plotted to show where in the auditory cortex each component explains the neural response. (B) Test-retest reliability of component responses to the natural sounds from this study. Each data point represents responses to a single sound, with color denoting its semantic category. Components 5 and 6 showed selectivity for speech and music, respectively, as expected (Component 4 also responded most to music due to its selectivity for sounds with pitch). (C) Component responses to natural and model-matched sounds constrained by the complete spectrotemporal model (see Figure S11 for results using subsets of model features). The speech and music-selective components show a weak response to model-matched sounds, even for sounds constrained by the full model. (D) Normalized squared error between responses to natural and model-matched sounds for each component.(E) The ratio of the standard deviation of each component’s responses to model-matched and natural sounds (see Figure S12A for corresponding whole-brain maps). (F) Pearson correlation of responses to natural and model-matched sounds (see Figure S12B for corresponding whole-brain maps). All of the metrics in panels D-F are noise corrected, although the effect of this correction is modest because the component responses are reliable (as is evident in panel B). Error bars correspond to one standard error computed via bootstrapping across subjects.

We found that all six components exhibited reliable responses to the natural sounds from this experiment (Figure 6B). Two of the components (5 & 6) responded selectively to speech and music, respectively, replicating the selectivity we found previously (last two columns of 6B). Critically, responses to the model-matched sounds were much weaker in these speech and music-selective components, even for sounds matched on the full model (Figure 6C, bottom row, last two columns; see Figure S11 for sounds matched on subsets of model-features), leading to high NSE values (speech NSE = 0.45; music NSE = 0.55 for the full model, noise corrected) (Figure 6D). By contrast, the other four components, all of which overlapped PAC to varying extents, responded similarly to natural and model-matched sounds constrained by the full model, leading to smaller errors (NSE < 0.27) than those for the speech and music selective components (p < 0.001 for all direct comparisons between the speech and music selective components and components 1, 2, and 4; for component 3, which had the lowest test-retest reliability, the direct comparison with the music-selective component was significant, p < 0.01, and the direct comparison with the speech-selective component was marginally significant, p = 0.076; statistics computed via bootstrapping across subjects). These results indicate that selectivity for music and speech cannot be purely explained by standard acoustic features that nonetheless account for much of the voxel response in primary regions.

Our model-matching approach posits that responses should be exactly matched if the model is accurate. If the model is not accurate, the approach makes no prediction about how the responses should differ. Nonetheless, the divergent responses to natural and model-matched sounds in Components 5 & 6 appeared to be largely driven by weaker responses to the model-matched sounds. We verified this observation by comparing the standard deviation of responses to natural and model-matched sounds: the response variation for model-matched sounds decreased sharply in Components 5 and 6 (Figure 6E). In contrast, the noise-corrected correlation remained high (Figure 6F). A similar pattern was also evident in whole-brain maps (Figure S12): the variation in voxel responses to model-matched sounds constrained by the full model dropped in non-primary regions (driven by lower responses to the “preferred” model-matched stimuli) while the correlation remained high. For Components 5 & 6, the high correlations were driven by the fact that model-matched sounds from the component’s preferred category produced a higher response than model-matched sounds from other categories (as is evident in Figure 6D). For example, in Component 6, model-matched music produced a lower response than natural music, but a higher response than model-matched sounds from other categories (p < 0.001; via bootstrapping). The same pattern was evident for Component 6, which responded selectively to speech (p < 0.001). This finding suggests that selectivity in non-primary regions may reflect a mixture of category-specific modulation tuning and responses to higher-order properties specific to music and speech, consistent with prior studies^53,60^. The results suggest that the modulation-specific structure driving Components 5 & 6 is correlated across natural sounds with the other properties of music and speech that drive their response. The model-matching approach allows us to see these two contributions to the response, revealing that there is something unique to the response of Components 5 & 6 that is distinct from the other components.

## Discussion

We have described a novel approach for evaluating a model of neuronal responses. Given a model, we synthesize a stimulus that yields the same model response as a natural stimulus, and test whether they produce similar neural responses. We applied this approach to test whether voxel responses in human auditory cortex can be explained by a commonly used auditory model based on spectrotemporal modulation. Our results revealed a substantial functional difference between primary and non-primary regions of human auditory cortex. Many voxels in PAC showed nearly equivalent responses to natural and model-matched sounds constrained by the full spectrotemporal model. We also found that these voxels responded differently when sounds were model-matched with only cochlear filter statistics, or with temporal or spectral modulations alone. These findings together suggest that spectrotemporal modulation accounts for much of the voxel response in PAC. By contrast, many voxels in non-primary regions responded weakly to all of the model-matched sounds, demonstrating that they are only weakly driven by the features captured by the model. This functional difference between primary and non-primary regions was not clearly evident when the model was evaluated by its response predictions, due to the confounding influences of stimulus-driven correlations across natural stimuli. Model-matching thus reveals a novel aspect of functional organization by showing where in the cortex a standard auditory model can explain voxel responses to natural sounds.

### Implications for models of auditory cortex

The notion that auditory cortex might be organized hierarchically – i.e., into a series of stages supporting increasingly abstract representations – has been a popular proposal for decades^40,61–63^. Hierarchical organization has some support from anatomical studies^64^, and from qualitative observations that responses in non-primary regions are more complex than those in primary regions^65,66^ and more closely aligned with semantically meaningful sound properties^16,45–47,67^. However, there has been little evidence for how primary and non-primary regions might differ in computational terms^41^, and thus it has been unclear what mechanisms underlie the apparent differences in tuning between primary and non-primary regions.

Most computational models of auditory processing beyond the periphery are based on tuning for modulation^19,20^. Such models have been used to explain responses throughout the auditory pathway in non-human animals^2,11,24–29,32,33^. In humans, such models have been shown to have relatively good predictive accuracy throughout both primary and non-primary regions^7,12,45^, which has led to the hypothesis that sounds are represented in a distributed manner^68^. This view contrasts with the notion of hierarchical organization, and in its most extreme form suggests that responses to seemingly complex attributes of sound in non-primary regions (e.g. speech and music-selectivity) could reflect the same types of mechanisms used to code sound in PAC.

Our study helps to reconcile these two prior literatures. First, we show that modulation selectivity fails to explain much of the response in non-primary regions, and that model predictions provide overly optimistic estimates of the model’s efficacy. This conclusion follows from the fact that we observed many voxels in non-primary regions whose response to natural sounds was well predicted by the model and yet produced divergent responses to model-matched sounds. Since the model by definition predicts that natural and model-matched sounds should have equivalent responses, this finding demonstrates a clear model failure. Since our predictions were based on linear regression, there must be some linear dependencies (i.e. correlations) across natural sounds between the model’s features and the other features which drive the voxel, and these correlations are what allow the model to accurately predict the voxel responses to natural sounds.

Conversely, our findings provide further evidence that modulation selectivity is a key feature of functional organization in human PAC^7,30,31^. Using both predictions and model-matching, we found that the modulation model explains the large majority of the voxel responses in this region. This finding was again not obvious from prior studies using model prediction alone, since the predictions could have been influenced by stimulus correlations, as turned out to be the case in non-primary regions. By contrast, we found that frequency selectivity, which presumably reflects tonotopy, explained much less response variance in PAC. Since fMRI is primarily sensitive to spatially clustered neural activity (because it pools responses from many nearby neurons), this finding suggests that modulation selectivity may be a key organizing dimension of PAC, and that neurons in auditory cortex with similar modulation selectivity may be spatially clustered.

What features might non-primary regions of auditory cortex represent? These regions are primarily driven by sound, show signs of having relatively short integration windows^43^, and even when speech selective, respond largely independently of the presence of linguistic structure^43,45^, suggesting acoustic rather than linguistic or semantic representations^16^. Moreover, although responses to the model-matched sounds were substantially weaker than responses to natural sounds, the model-matched sounds still drove responses to natural sounds above baseline and were correlated with responses to natural sounds. Thus, one natural hypothesis is that non-primary regions transform a lower-level acoustic representation like the spectrotemporal representation considered here, into an acoustic representation that makes behaviorally relevant variables more explicit (e.g. easier to decode). We have recently explored this idea by training a deep neural network to recognize words and musical genres^41^ and then comparing the resulting representations to voxel responses. We found that later layers of the network better predicted voxels in non-primary regions of the cortex, consistent with the notion of hierarchical organization. These predictions could of course be influenced by stimulus-driven correlations, which may explain why the differences in prediction accuracy between layers were modest. Future work could address this question by applying model-matching to the representation from different layers of a hierarchical model.

### Implications and limitations of model-matching

The result of our model-matching experiment is an error metric between 0 and 1 indicating the dissimilarity of a neural response to natural and model-matched sounds. What does this number tell us about the type of models that could underlie the neural response? When the error metric is near 1, the models under which responses have been matched are ruled out as descriptions of the neural response underlying the voxel. Our specific implementation matched model responses for all point-wise functions of the filters in question, and thus that family of models is ruled out for voxels with large error.

At the other extreme, errors near 0, like those we observed in primary auditory cortex, reveal that the voxel responses are consistent with the family of models whose response was matched. The matching procedure employed a specific filter bank, but alternative models might also be matched (for instance those with filters that can be approximated as linear combinations of the filters used for matching). Small error values thus do not exclude models other than the one we used. However, specific alternative models could be evaluated by measuring their response to the two sets of stimuli used in an experiment (natural and model-matched). Models that give distinct responses to the two stimulus sets could be ruled out for voxels whose responses to the two sets are similar. Conversely, one could also rule out models whose responses to the two sets are similar for voxels whose responses to the two sets are different. We used this approach to investigate different types of spectrotemporal filter banks (Figure S15), finding that a range of alternative filter banks had matched statistics for the natural and model-matched sounds tested here (see *Variants of the spectrotemporal filter model* in the Methods). This finding suggests that a wide range of spectrotemporal filter models can be ruled out as models of non-primary auditory cortex. Our stimuli and neural data will be made available online upon publication, so that alternative models can be evaluated using this approach.

In other situations, matching with one model may entail matching with another, but not vice versa. This was the case for the four models we compared in Figure 4 – the full spectrotemporal model is inclusive of the other models. The weaker correlations observed with the other models provides evidence for the necessity of the spectrotemporal model features.

As with any method, the interpretation of our results is constrained by the resolution of the neural measurements. Since fMRI is primarily sensitive to brain responses that are spatially clustered, our results bear most directly on aspects of cortical tuning that are organized at a relatively large spatial scale across the cortex. Our results were robust to the exact size of the voxels tested and the amount of spatial smoothing, suggesting that our results hold for spatial scales on the order of millimeters to centimeters. But even small voxels pool activity across neurons and across time, and thus it is possible that voxels with similar responses to natural and model-matched sounds might nonetheless contain neurons that show more divergent responses, or that have temporal response properties that differ from the model. This fact may partially explain why electrophysiological recordings in animals have found that linear spectrotemporal filters are insufficient to account for responses in primary auditory cortex^13,69–71^.

By contrast, the implication of divergent voxel responses is more straightforward – they imply that the underlying brain responses must also differ. In principle it is possible that averaging across neurons could average out neural responses that are heterogeneous across neurons but similar between responses to natural and model-matched sounds, making voxel responses to the two sets of stimuli more dissimilar than in the underlying neural populations. However, the difference in voxel responses between natural and model-matched sounds evident in our experiments was largely due to substantially lower responses to model-matched sounds, suggesting that the firing rate of neurons within the voxel was also lower, demonstrating that the model lacks features which drive the response.

Future work could apply model-matching to neuronal responses measured electrophysiologically to test models at a finer spatial and temporal scale. For example, one could synthesize model-matched sounds that should yield the same firing rate as a natural sound given a model of an individual neuron’s response.

### Relation to prior work on perceptual metamers and texture synthesis

Our approach to model matching is an extension of methods for texture synthesis originally developed in image processing and computer vision^48,72^, and later applied to sound texture^39^ and visual texture perception^49,73^. In texture synthesis, the goal is typically to test whether a set of statistical features could underlie perception, by testing whether synthetic stimuli with the same statistics are metameric, i.e. whether they look or sound the same as a real-world texture. The implementation of our synthesis procedure is inspired by classic texture synthesis methods^48^, but the scientific application differs notably in that we evaluate the model by the similarity of neural responses rather than the similarity that is perceived by a human observer. Indeed, many of the model-matched stimuli sounded unnatural, which demonstrates that the modulation spectrum fails to capture higher-order properties of natural sounds that listeners are sensitive to (e.g. the presence of phonemic or melodic structure). This observation demonstrates the insufficiency of the modulation spectrum as a complete account of perception, but not does place strong constraints on which neural stages are well described by the model. The fact that responses to natural and model-matched sounds diverged in non-primary regions of auditory cortex suggests that they may be driven by higher-order structure not made explicit by the modulation model.

The most similar previous approach involved comparing the strength of cortical responses to visual textures synthesized from different classes of statistics of a wavelet filter bank model^74^. Although we also compared cortical responses to sounds synthesized from different model statistics, the key comparison was between responses to individual natural and synthesized sounds, which is critical to identifying regions of the brain that are not well explained by a model.

The modulation filter bank model tested here bears similarities to the texture model described in McDermott & Simoncelli (2011). The key difference is that dependencies between cochlear frequency channels were captured by spectral modulation filters rather than the correlations used in the original texture model. In practice, we found that sounds synthesized from the two models were perceptually similar, suggesting that correlations in one stage of representation (the cochlea) can be captured by marginal statistics of a subsequent stage of representation (modulation filters), as noted by McDermott & Simoncelli.

### Approaches for model testing

Recent years have seen growing interest in the use of computational “encoding models” to test formal theories of sensory processing^5–7,10–14,16,17,75,76^. Because encoding models make quantitative predictions about the neural response, they can be used to test and compare theories of neural coding. The features of the model can then provide insight into the sensory features that are represented in different neural populations^6,7,12,16^.

A key challenge of testing encoding models with natural stimuli is that the features of different models are often correlated^16,18^, making it difficult to tease apart the unique contribution of any particular model. This problem can be partially overcome by comparing the predictions of two different models, but is difficult to eliminate when the features of two models are strongly correlated and when responses can only be measured to a relatively small number of stimuli (as is common with fMRI). Another approach is to alter stimuli so as to decouple different features sets^76^. For example, adding varied background noise to natural sounds could help to decouple low and high-level features of sounds, because noise can alter a sound’s low-level features without affecting its perceived identity. However, such approaches are heuristic, and do not guarantee that the relevant features will be de-correlated unless the candidate feature sets can be measured with existing models. Model-matching is appealing because it provides a way to test the ability of a single model to explain neural responses by imposing the structure of that model alone, decoupling the model from alternative models without needing to specify the many possible alternatives.

## Author Contributions

SNH and JHM designed the stimuli and experiment together and wrote the paper. SNH implemented all of the experimental procedures, synthesis algorithms, and analyses.

## Methods

### Participants

The experiment comprised 41 scanning sessions, each approximately two hours. Fifteen subjects participated in the experiment (ages 19-36; 5 male; all right-handed; one subject, S1, was author SNH). Two different experiment paradigms were tested (hereafter referred to as Paradigm I and Paradigm II). We have chosen to describe these two paradigms as a part of the same experiment because the stimuli and analyses were very similar. In Paradigm I, eight subjects completed a single scanning session, three subjects completed five sessions, and one subject completed three sessions (this subject chose not to return for the 4^th^and 5^th^sessions). We chose this approach because it allowed us to compute reliable group maps by averaging across the twelve subjects, as well as reliable individual subject maps using a larger amount of data from the subjects with multiple scan sessions. Five subjects were scanned in Paradigm II. One subject completed two sessions, two subjects completed three sessions, and one subject completed four sessions. One subject (S1) was scanned in both paradigms (when possible we used data from Paradigm II for this subject, because there was a higher quantity of data, and the scan sessions for Paradigm II were higher resolution as noted below).

Because we aimed to characterize the auditory cortex of typical listeners without extensive musical experience, we required that subjects not have received formal musical training in the five years preceding their participation in the experiment. The study was approved by MIT’s Committee on the Use of Humans as Experimental Subjects; all subjects gave informed consent.

### Data acquisition parameters and preprocessing

Data for Paradigm I were collected on a 3T Siemens Trio scanner with a 32-channel head coil (at the Athinoula A. Martinos Imaging Center of the McGovern Institute for Brain Research at MIT). The functional volumes were designed to provide good spatial resolution in auditory cortex. Each functional volume (i.e. a single 3D image for one participant) included 15 slices oriented parallel to the superior temporal plane and covering the portion of the temporal lobe superior to and including the superior temporal sulcus (3.4 s TR, 30 ms TE, 90 degree flip angle; 5 discarded initial acquisitions). Each slice was 4 mm thick and had an in-plane resolution of 2.1 × 2.1 mm (96 × 96 matrix, 0.4 mm slice gap). iPAT was used to minimize acquisition time (1 sec/volume). T1-weighted anatomical images were also collected for each subject (1 mm isotropic voxels).

Data for Paradigm II were collected more recently using a 3T Prisma scanner (also at the McGovern Institute). We used a multiband acquisition sequence (3x acceleration) to reduce slice thickness while maintaining coverage (36 slices with 2mm thickness and no gap), and thus reducing voxel size (2 mm isotropic). iPAT was not used. Other acquisition parameters were similar (3.45 s TR, 1.05 sec acquisition time, 34 ms TE, 90 degree flip angle; 3 discarded initial acquisitions).

Functional volumes were preprocessed using FSL software and custom MATLAB scripts. Volumes were motion-corrected, slice-time-corrected, skull-stripped, linearly detrended, and aligned to the anatomical volumes (using FLIRT^77^ and BBRegister^78^). Volume data were then resampled to the reconstructed cortical surface computed by FreeSurfer^79^, and smoothed on the surface using a 5mm FWHM kernel to improve SNR (results were similar without smoothing; Figure S3). Individual subject data were then aligned on the cortical surface to the FsAverage template brain distributed by Freesurfer.

### Stimulus presentation and scanning procedure

Our stimulus set was derived from 36 natural sounds, each 10-seconds in duration (Figure 2B). From each natural sound, we synthesized four model-matched sounds, constrained by different subsets of features from a model of auditory cortex based on modulation filtering^19^. The complete stimulus set thus included 5 conditions (natural sounds + 4 model-matched versions) each with 36 sounds, yielding a total of 180 stimuli.

Scan acquisitions produce a loud noise due to rapid gradient switching. To prevent these noises from interfering with subjects’ ability to hear the sounds, we used a “sparse” scanning paradigm^80^that alternated between presenting sounds and acquiring scans, similar to those used in our prior experiments^45,81,82^(Figure S2). This was achieved by dividing each 10-second stimulus into five 2-second segments (windowed with 25 ms linear ramps). These five segments were presented sequentially with a single scan acquired after each segment. The five segments for a particular sound were always presented together in a “block” (the order of the segments within a block was random). Each scan acquisition lasted 1 second in Paradigm I and 1.05 seconds in Paradigm II. There was a 200 ms buffer of silence before and after each acquisition. The total duration of each five-segment block was 17 seconds in Paradigm I and 17.25 seconds in Paradigm II. We averaged the response of the 2^nd^ through 5^th^ acquisition after the onset of each stimulus block. The first acquisition was discarded to account for the hemodynamic delay. Results were similar when we instead averaged just the second and third timepoint or just the fourth and fifth timepoint after stimulus onset, indicating that our results were robust to the averaging window applied to the fMRI measurements (Figure S13). We chose to use signal averaging rather than a GLM with a standard hemodynamic response function (HRF), because we have found this approach is leads to slightly more reliable responses, presumably due to inaccuracies in the standard HRF^83^.

In Paradigm I, each model-matched stimulus was presented once per two-hour scanning session, and the natural stimuli were presented twice so that we could measure the reliability of each voxel’s response to natural sounds and noise-correct the normalized squared error metric. Each session was divided into 12 “runs”, after which, subjects were given a short break (∼30 seconds). Each run included 6 natural sounds and 12 model-matched sounds (3 per condition). In Paradigm II, we presented only the model-matched sounds constrained by the complete model, which allowed us to present both the natural and model-matched sounds several times per scan session. Each run included 9 natural and 9 model-matched sounds. The entire sound set was presented over 4 consecutive runs. Subjects completed 12 or 16 runs depending on the time constraints of the scan session. Thus, each subject heard each sound between 3 or 4 times per session. In both paradigms, there were periods during which no stimulus was presented and only scanner noise was heard, which provided a baseline with which to compare stimulus-driven responses. There were four such “silence” periods per run (each 17 seconds in Paradigm I and 17.25 seconds in Paradigm II). The ordering of stimuli and silence periods was pseudorandom, and was designed such that on average each condition occurred with roughly the same frequency at each position in a run, and each condition was preceded equally often by every other condition (as in our prior work^81,82^).

Prior to settling on the procedure for Paradigm I, we conducted a pilot experiment in which six of the twelve participants from Paradigm I completed a single session. These sessions featured stimuli from only 3 of the model-matched conditions (spectral modulation matched stimuli were omitted). These scan sessions were the first of this study and we limited the number of conditions to make sure the experiment could fit within the allotted 2-hour scanning slot. The runs for these sessions were slightly shorter because there were only 9 model-matched stimuli presented per run (there were only 3 periods of silence per run for these sessions). When analyzing the results we included the data from these sessions in order to use the maximum amount of data available for each condition, and thus the results for the spectral modulation matched condition were based on less data than the other model-matched conditions. But because the NSE metric was corrected for noise (see below), differences in the amounts of data across conditions should not bias the results.

### Selection of natural stimuli

We used long sounds (10 seconds) so that we could compute time-averaged statistics for filters with relatively long integration periods (i.e. periods of up to 2 seconds). We selected sounds that were likely to produce high response variance in auditory cortical voxels, guided by the results of a prior paper from our lab that measured fMRI responses in auditory cortex to a large set of natural sounds^45^. In our prior study, we found that much of the response variance could be captured by a weighted sum of six response patterns (“components”), and we thus attempted to select sounds that had high response variance along these components. To accomplish this goal, we created a subset of 60 sounds with high component response variance by iteratively discarding sounds in a greedy manner, each time removing the sound that led to the largest increase in response variance averaged across the six components. Because we needed stimuli that were relatively long in duration, we could not directly use the stimuli from our prior study, which were only 2-seconds in duration. Instead, we created a new stimulus set with 10-second sounds, each of which had the same label (e.g. “finger tapping”) as one of the sounds from the 60-sound set.

### Model representation

We synthesized sounds based on four different model representations. The simplest model was just based on the output of filters designed to mimic cochlear responses (i.e. a cochleagram). The other three models were based on filters tuned to modulations in this cochleagram representation. Two models were tuned to either temporal modulation or spectral modulation alone, and one was jointly tuned to both temporal and spectral modulation.

The cochlear representation was computed by convolving the audio waveform of each sound with 120 bandpass filters, spaced equally on an ERB_N_-scale between 20 Hz and 10 kHz, with bandwidths chosen to match those measured psychophysically in humans (individual filters had frequency responses that were a half-cycle of the cosine function in order to exactly tile the frequency spectrum; adjacent filters overlapped by 87.5%)^39^. Each channel was intended to model the response of a different point along the basilar membrane. The envelopes of each filter output were computed using the Hilbert transform, raised to the 0.3 power to mimic cochlear compression/amplification, and downsampled to 400 Hz after applying an anti-aliasing filter. So that we could express the spectral modulation filters that operate on the cochleagram (described below) in units of cycles per octave (as in Chi et al., 2005^19^, we interpolated the frequency axis from an ERB-scale to a logarithmic frequency scale (24 cycles/octave), yielding 217 channels.

The modulation-based representations were computed using a bank of multi-scale wavelet filters (Figure 1A). The shapes and bandwidths of the filters were the same as those described by Chi et al. (2005). The three sets of filters differed in whether they were tuned to modulation in time, frequency or both. The temporal modulation representation was computed using gammatone filters:

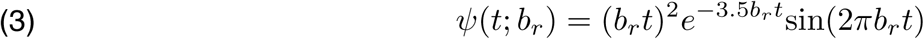

where *b*_*r*_ determines the best modulation rate of the filter (i.e. the rate with maximum gain). We used 9 filters with octave-spaced best rates: 0.5, 1, 2, 4, 8, 16, 32, 64, and 128 Hz. Each filter was separately convolved in time with each frequency channel of the cochleagram. The output of the model can thus be represented as a set of 9 filtered cochleagrams, each of which highlights modulations at a particular temporal rate.

The spectral modulation representation was computed using ‘mexican hat’ filters, which are proportional to the second derivative of a Gaussian:

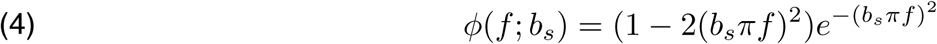

where *b*_*s*_ determines the best modulation scale of the filter (i.e. the scale with maximum gain). We used 6 filters with octave-spaced scales: 0.25, 0.5, 1, 2, 4, and 8 cycles/octave. Each filter was separately convolved in frequency with each time “slice” of the cochleagram. The output of the model can thus be represented as 6 filtered cochleagrams, each of which highlights a different range of spectral modulations.

The spectrotemporal modulation representation (often referred to as the “full model”) was computed primarily from 2D filters that were convolved with the cochleagram in both time and frequency. The filters were instantiated in the 2D Fourier domain (as described in Chi et al., 2005^19^) by taking the outer-product of the frequency-domain representations of the temporal and spectral modulation filters described above. These filters were then ‘oriented’ so as to be sensitive to upward-right or downward-right modulations. This was accomplished by zeroing either the first and third quadrant of the 2D frequency response (for upward-oriented filters), or the second and fourth quadrant (for downward-oriented filters) (the Nyquist frequency and DC were never zeroed). There were 108 total spectrotemporal filters produced by crossing 9 temporal filters with 6 spectral filters (with best modulation frequencies as described above), and orienting each filter upwards or downwards. Thus, the output of this portion of the model can be represented by 108 filtered cochleagrams (modulo the additional filters described next).

For all three modulation-based representations (temporal, spectral and spectrotemporal), we included the unfiltered cochleagrams in the representation so that the modulation-based representations would be strictly more expressive. For both the temporal and spectral representations we also included a filter with power at only the DC (0 Hz or 0 cycles/octaves respectively). These filters capture the mean of each cochlear frequency channel (for the temporal modulation representation) or the mean of each time slice through the cochleagram (for the spectral modulation representation), and were necessary to reconstruct cochleagrams from the model representation (because all of the other filters were bandpass, with zero power at the DC). For the spectrotemporal modulation representation, the temporal and spectral DC filters were also crossed with the other filters, yielding an additional 15 filters; these filters capture spectrally broadband temporal modulations (i.e. “vertical” modulations), or temporally uniform spectral modulations (i.e. “horizontal” modulations) and have only one orientation. We also added all of the filters from the temporal-only and spectral-only modulation models to the spectrotemporal modulation model so that it would be strictly more expressive than the simpler models.

Finally, two filters which were modulated only in time and which had very low best-modulation rates (0.125 and 0.25 Hz) were added to the temporal and spectrotemporal modulation representations. These filters were included to replicate the homogeneity of the natural sounds in the model-matched sounds and to improve convergence. Without them, the synthesis process tended to “clump” sound energy at particular time-points. The low-rate filters ameliorated this problem by forcing the slow fluctuations in the model-matched sounds to be similar to those in the natural sounds they were matched to.

### Model-matching synthesis algorithm

Our model-matching approach, like most algorithms for texture synthesis, starts with a sample of noise, which initially lacks structure, and alters the noise via an iterative procedure to match statistical constraints^39,48,72^, in our case provided by the histogram of each feature’s response (Figure S1). By initializing with noise, we aim to arrive at a sound that is minimally structured given the imposed constraints.

The model-matching synthesis procedure was initialized with a 10-second sample of Gaussian noise (the same duration as the natural sounds). The algorithm involved three steps: (1) computing the response of each feature from a given model to a natural and noise sound (2) separately matching the response histogram across time for each model feature^48^ (3) reconstructing a waveform from the modified outputs. These three steps were applied iteratively for reasons described below. For the cochlear representation, we matched the histogram of envelope values for each cochlear frequency channel. For the modulation-based representations, we matched the histogram of each frequency channel of each of the filtered cochleagrams (each channel of the filtered cochleagrams represents the output of a single model feature), as well as the histograms of the unfiltered cochleagram frequency channels.

The goal of our histogram matching procedure was to modify the distribution of values for one time series so that they had the same distribution of values as that of a target time series, without imposing the same temporal pattern over time. For example, to modify the time series [1 2 3] to match the histogram of [5 1 3], we would like to alter the first time series to be [1 3 5], such that it has the same distribution as the target, but the same relative ordering as the original (smallest, middle, largest). Since the average value of a signal only depends on the distribution of magnitudes and not their ordering, the histogram-matched signals will have the same average value, even if they are transformed by a point-wise function (e.g. 1^2^+ 3^2^ + 5^2^ = 5^2^+ 1^2^+ 3^2^). Assuming the two time series are represented as vectors of equal length, as was the case for our experiments (because the synthetics were of equal duration), we can histogram match the signals by re-assigning the smallest value in the signal-to-be-matched to the smallest value in the target signal, then re-assigning the second smallest value in the signal-to-be-matched to the second smallest value in the target, and so on. We can implement this procedure with the following pseudocode:

order_original = sortindex(original)
order_target = sortindex(target)
matched[order_original] = target[order_target]

where sortindex is a function that takes a vector as input and returns a list of indices into that vector which have been ordered according to the magnitude of the corresponding vector elements (i.e. the indices that would sort the vector, as in the numpy/python function argsort). This procedure is a slightly simpler variant of the histogram-matching algorithm described by Heeger and Bergen (1995), and is applicable when matching vectors of equal length.

The details of the reconstruction algorithms have been described previously^19,39^. We reconstruct a waveform from a cochleagram by summing the individual subbands, which are computed by multiplying the envelopes of each cochlear channel (after histogram matching) by their time-varying phases from the previous iteration and then refiltering with the filters used to generate the subbands (as is standard for subband transforms). Similarly, we reconstruct a cochleagram from the modulation domain by adding up the filtered cochleagrams in the 2D Fourier domain and multiplying each cochleagram by the complex conjugate of the filter (to undo phase shifts). We then divide by the summed power of the modulation filters to correct for the fact that the summed power of the filters is not uniform^19^.

Because the filters whose outputs are being manipulated overlap in the frequency domain, a manipulation such as histogram matching typically introduces inconsistencies between filters, such that when the reconstructed signal is re-analyzed the histograms will generally not remain matched. As such, a single iteration of the matching procedure does not achieve its objective, but iterating the procedure (analyze, histogram-match, reconstruct) generally results in increasingly close matches^39,48,72^. We monitored convergence by measuring the difference in the desired and measured histograms at each iteration (see below), and used 100 iterations, which we found to produce good results.

To avoid wrap-around effects due to circular convolution, we padded the cochleagrams in time and frequency prior to convolution with the modulation filters. We padded the cochleagrams with a value equal to the global mean of the cochleagram across both time and frequency so as to minimize the resulting step edge. The amount of padding was chosen to be approximately equal to the duration of ringing in each filter: we padded the cochleagrams in frequency by twice the period of the coarsest spectral modulation filter (8 octaves of padding) and by three times the period of the slowest temporal modulation filter (24 seconds of padding). To ensure that the portion of the signal used for the stimulus was well matched, we applied the histogram matching procedure twice at each iteration, once to the entire signal including the padded duration, and once to just the non-padded portion of the signal.

MATLAB code for synthesis algorithm will be made available upon publication.

### Assessing the success of the model-matching algorithm

For each model feature, we computed a time-averaged measure of its response amplitude for natural and model-matched sounds. Figure S14 plots these amplitude statistics for example natural and model-matched sounds. For cochlear features, we simply averaged the cochleagram envelope amplitudes across time. For the modulation-tuned features, we computed the standard deviation across time of each feature’s response. We then correlated the filter amplitudes for corresponding natural and model-matched sounds across all filters in the model, as a measure of their similarity (Figure S14, right panel). The mean correlation across sounds was high for all of the model features being matched by the synthesis algorithm (r^2^ > 0.98), and much higher than the correlation observed for features not constrained by the matching algorithm.

### Variants of the spectrotemporal filter model

We investigated the extent to which our results might depend on the particular choice of spectrotemporal filters tested (Figure S15). Specifically, we created spectrotemporal filters with different properties by either (1) randomizing the temporal and spectral phase to create a diverse range of filter shapes with roughly the same modulation spectrum (2) halving the filter bandwidths or (3) randomizing the filters entirely by sampling the filter weights from a Gaussian. Phase randomization was implemented by computing the FFT of each filter’s temporal and spectral impulse response (using a window size of twice the period of the filter’s center modulation rate/scale), randomizing the phase, transforming back to the signal domain (via the iFFT), and padding with zeros. Narrowing the filter bandwidths was accomplished by halving the temporal and spectral extent of the filters in the frequency domain as well as doubling the number of filters to ensure that all modulation rates and scales were represented by the model. For the random filters, we varied the size of the filters to mimic the fact that the model filters vary in the amount of time and frequency over which they integrate.

For each filter, we measured the amplitude (standard deviation) of its response to each of the natural and model-matched sounds that we tested in the fMRI experiment (middle panels of Figure S15), which were constrained only by the original spectrotemporal filters and not the modified variants. We then correlated the filter’s amplitude for corresponding natural and model-matched sounds (across the bank of filters for each sound) to assess how well the natural and model-matched sounds were matched (rightmost panel of Figure S15). For the phase-randomized and half-bandwidth filters, we found that matching the spectrotemporal statistics of the original filters substantially improved how well the modified spectrotemporal filters were matched (median r^2^ > 0.85), suggesting that matching the statistics of one spectrotemporal model goes a long way toward matching the statistics of other modulation filter models. This result suggests that our findings will generalize to other spectrotemporal filters with different shapes. For random filters, we found that the filter variances were relatively well matched even for sounds that were not matched on the original spectrotemporal filters (median r^2^ = 0.73 for cochlear matched sounds), suggesting that random filter variances may be easier to match than the more structured filters in the model.

### Envelope-based synthesis algorithm

The stimuli used for the first six pilot scan sessions were synthesized using a slightly different algorithm that was based on matching the histogram of the envelopes of the modulation filter outputs rather than matching the histogram of the raw filter responses. In practice, we found that histogram matching the envelopes produced very similar results to matching the histogram of the raw outputs, and thus decided to use the simpler algorithm for the remaining scanning sessions. The voxel responses to stimuli synthesized from the two algorithms was similar, and we thus collapsed across all of the available data for all analyses.

### Normalized squared error

We measured the similarity of fMRI responses to natural and model-matched sounds via the mean squared error:

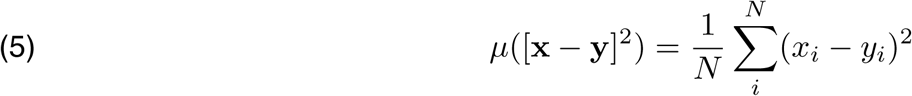

where **x** and **y** represent the vector of responses to natural and model matched sounds, respectively (here N=36 because there were 36 natural/model-matched sounds). We normalized the mean squared error so that it would be invariant to the overall scale of the voxel responses and take a value of 0 if the response to natural and model-matched sounds was identical and 1 if there was no correspondence between responses to natural and model-matched sounds (i.e. if they were independent of each other):

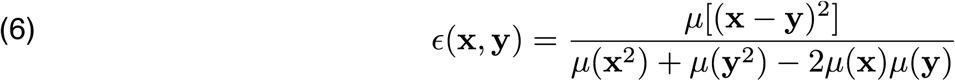

We refer to this metric as the normalized squared error or NSE. The quantity in the denominator is an estimate of the expected value of the squared error assuming the two variables are independent:

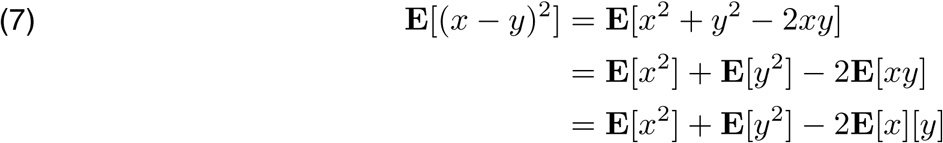

### Noise-correcting the normalized squared error

Because fMRI responses are noisy, the error between two fMRI measurements or between a measurement and a prediction will be typically be higher than the “true” error in the absence of noise. Thus, we chose to correct the NSE for measurement noise to ensure that differences between voxels were not driven by differences in voxel noise, and to provide an estimate for what the true NSE would be in the absence of noise. In practice, we observed similar trends with and without correction because voxels responses in both primary and non-primary regions were similarly reliable (Figure S4).

Most noise-correction methods assume that the noise-corrupted response reflects the sum of a noise-free stimulus-driven signal plus noise that is statistically independent of the stimulus-driven signal:

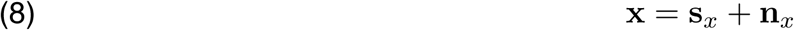

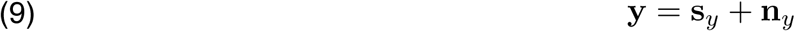

where in the context of this experiment, **x** and **y** are the measured response of a voxel to two sets of sounds (i.e. natural and model-matched sounds), **s***x* and **s***y* are the stimulus-driven responses, and **n***x* and **n***y* are the noise that contributes to the response measurements. All noise-correction methods require at least two repetitions of the same stimulus so that the effects of the noise can be disentangled from the effects of the stimulus-driven signal. By assumption these two repetitions only differ in their noise:

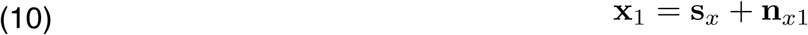

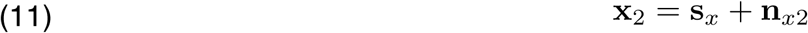

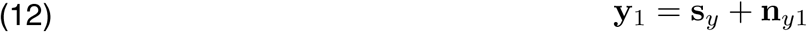

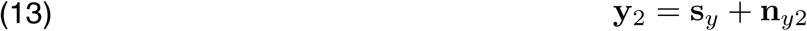

We would like to estimate the NSE of the stimulus-driven component of the uncorrupted responses:

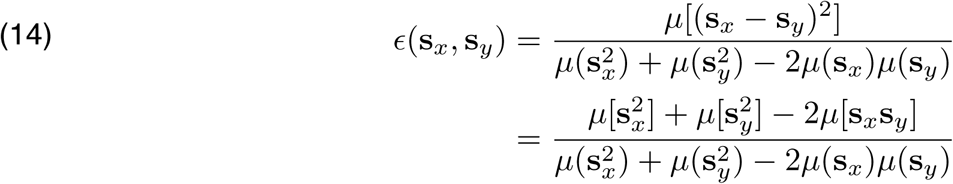

But are restricted to measuring the NSE of the noise-corrupted response. From the equation above, it is evident that the NSE depends on three types of statistics: (1) the signal powers (u[s_x_^2^] and u[s_y_^2^]) (2) the signal cross-product (u[s_x_s_y_]) and (3) the signal means (u[s_x_]). The signal means are unbiased by the noise, since by assumption the noise is zero mean. The signal cross-product is also unbiased by noise:

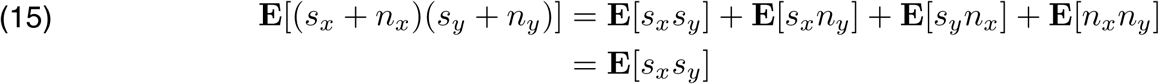

(we have replaced means with expectations to indicate a theoretical average over infinitely many samples, for which the bias is exactly zero). We thus estimate the signal cross-product and means using the measured cross-product and means of the data without correction:

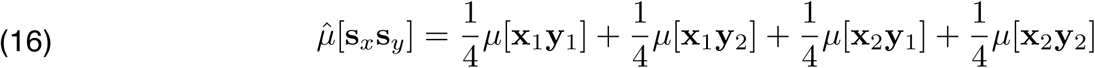

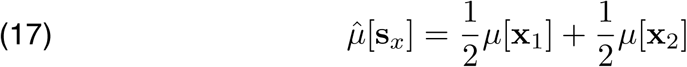

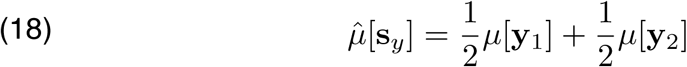

Unlike the mean and the cross-product, the signal power is biased upwards by the noise:

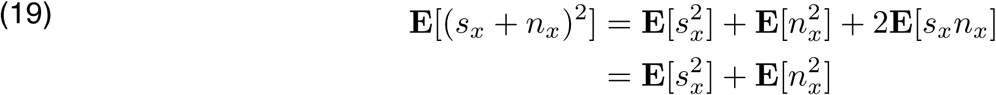

The magnitude of this bias can be estimated using the residual error between two measurements of the same stimulus, which by definition is due exclusively to noise. The expected power of the residual is equal to twice the noise power:

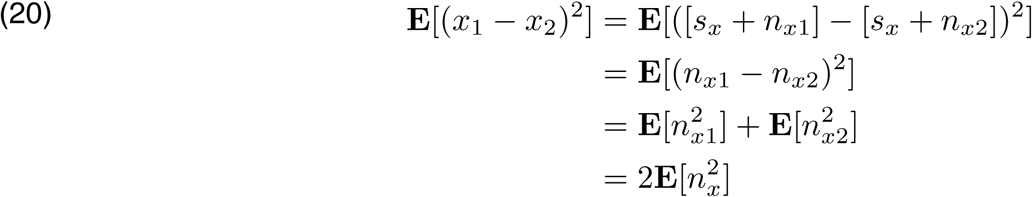

Thus, we can estimate the signal power by subtracting off half the residual power from the average power of the noise-corrupted data:

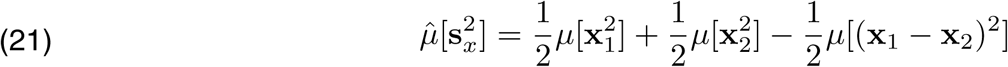

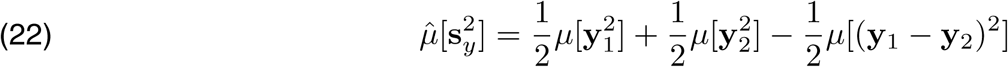

Substituting equations 16-18 and 21-22 into equation 14 yields the noise-corrected NSE. The noise-corrected NSE, like the raw NSE, is invariant to the overall scale of the data.

Noise-correction requires two independent samples of the same stimulus. In our case, each sample was itself an average across multiple stimulus blocks, and for each stimulus block, we averaged responses across the last four scan acquisitions within the block. Thus, each sample was based on many scan acquisitions (between 12 and 28 acquisitions for individual subject maps, corresponding to between 3 and 7 stimulus block repetitions; group maps were based on 104 scan acquisitions per measurement). In Paradigm I, each natural sound was repeated once per scan while the model-matched sounds were only presented once. We chose this design so that we could present model-matched sounds constrained by different subsets of model features, which would have been infeasible if each model-matched sound was presented twice. To noise-correct the responses, we made the simplifying assumption that the noise power was equal for natural and model-matched sounds, and estimated the noise power from responses to the natural sounds (when multiple scan sessions were available we first averaged responses across scan sessions). This assumption is natural given that the noise by definition reflects the component of the signal that is not driven by the stimulus. Nonetheless, we tested whether this assumption is appropriate using the data for Paradigm II, in which we repeated responses to both natural and model-matched sounds. In one case, we assumed that the noise power was the same, and calculated the noise power using only the responses to natural sounds. In the other case, we separately calculated the noise power for natural and model-matched sounds. The results were very similar using the two approaches (Figure S16), which validates the assumption that the noise power is similar for natural and model-matched sounds.

Our noise correction procedure assumes that the noise is uncorrelated across measurements (this assumption was used in equation 15), which is the not the case for fMRI measurements close in time (i.e. <5 seconds)^84^. Here, each measurement corresponds to the average response of the 2^nd^ through 5^th^ scan acquisition after the onset of each stimulus block. Blocks for the same stimulus were never repeated back to back, and even if they were, the two blocks would have been separated by 6.8 seconds, which is longer than the typical autocorrelation of the BOLD signal^84^. In Paradigm II, the same stimuli were never repeated within a run. Thus, it is unlikely that the autocorrelation of the BOLD signal impacted our measures.

### Evaluating the noise-corrected NSE with simulated data

Noise-correction inevitably increases the variance of the statistics being corrected, and thus it is critical to have sufficiently reliable responses (which is why we collected a relatively large amount of data for this study). To assess the reliability needed to perform correction, we performed a simulation in which we generated a large number of noisy voxel responses. We based our simulations on Paradigm I in which only the natural sounds were repeated, but results were similar for simulations that mimicked Paradigm II where both natural and model-matched sounds were repeated. For Paradigm I, we had three 36-dimensional response vectors per voxel: two vectors for the 36 natural sounds, which were each presented twice per scan session, and one for the model-matched sounds, which were each presented once per scan session. We thus simulated three 36-dimensional response vectors (**x**_1_, **x**_2_, **y**_1_) for each voxel (**x**_1_ and **x**_2_ corresponding to the voxel’s response to natural sounds, and **y**_1_ to the response to model-matched sounds). Each vector was computed as the weighted combination of a true, noise-free signal (**s**_x_, **s**_y_) that was constant across repeated measurements plus additive noise that varied across measurements (**n**_x1_**, n**_x2_, **n**_y1_):

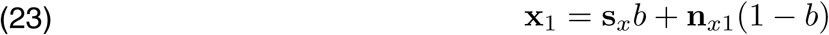

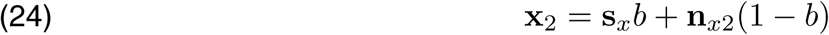

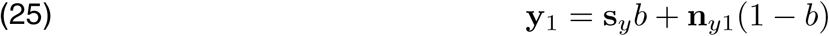

We used the weights (*b*) to control the SNR of the voxel with weights closer to 1 resulting in higher SNR. We sampled the three noise vectors from a zero-mean, unit-variance Gaussian distribution. Our noise-correction algorithms assume that the noise variance is the same for the natural and model matched sounds (var(n_x_) = var(n_y_)), which we have verified is a reasonable assumption for our data (Figure S16). We also assume that the noise samples are independent from each other, which we would expect to be the case given that our measurements were spaced far apart in time relative to the autocorrelation of the BOLD signal^84^. Our noise-correction algorithm makes no assumptions about the distribution of errors. Here we use a Gaussian distribution for simplicity, but results were similar using other noise distributions (e.g. Laplace).

We sampled the true noise-free signals (**s**_x_ and **s**_y_), in a way that allowed us to vary how similar they were. We did this in two different ways (referred to hereafter as Simulation 1 and Simulation 2). In Simulation 1, we computed **s**_x_ and **s**_y_ as the weighted sum of a shared response vector (**g**) and a distinct vector unique to x and y (**u**_x_, **u**_y_):

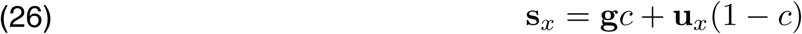

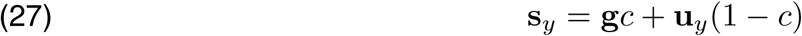

Thus by varying c, we could vary the similarity of the voxel responses. We sampled c from a uniform distribution, and we sampled g, u_x_, and u_y_ from a zero-mean, unit-variance Gaussian. The results were similar using other distributions (e.g. the Gamma distribution). Changing the means of these distributions also had little effect on the results.

In Simulation 2, one of the signal vectors was simply a scaled version of the other, in order to mimic weaker responses to model-matched sounds:

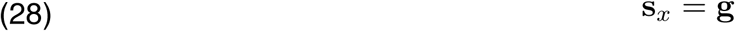

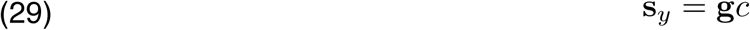

For each type of simulation, we sampled 100,000 voxel responses. For each sample, we computed four statistics:

1. the NSE between the noisy signals (using just **x**_1_ and **y**_1_ for simplicity).
2. the NSE between the true signals (**s**_x_ and **s**_y_), which is what we would like to infer.
3. our estimate of the true NSE, computed by applying our noise-correction algorithm to the noisy data (**x**_1_, **x**_2_ and **y**_1_).
4. the NSE between two independent measurements of the same stimulus (“test-retest”), which provides a measure of the voxel’s noise level (**x**_1_, **x**_2_).

In Figure S17A&B, we plot the results from Simulation 1. First, we plot the NSE of the noise-corrupted data vs. the NSE of the true signals (Figure S17A, left column). Each point represents the NSE values for a single simulated voxel response, and the results have been binned by the test-retest NSE values of the noise-corrupted signals (from low to high going from top to bottom of the figure), which provides a measure of the noise level (lower test-retest NSEs corresponding to less noise). Unsurprisingly, as the noise increases, the upwards bias caused by the noise increases. Next, we plot the noise-corrected NSE values vs. the NSE values for the true signals (Figure S17A, right column). As expected, noise-correction removes the bias caused by the noise, at the expense of increasing the variance. These effects are quantified in Figure S17B, which plots the median NSE of both the noise-corrupted and noise-corrected values along with its standard deviation (central 68% of the sampling distribution). At high noise levels (test-retest NSE>0.4), noise-correction substantially increases the standard deviation of the samples, which makes correction untenable. But for low noise levels (test-retest NSE <0.4), the method corrects the bias without substantially increasing the standard deviation of the sampling distribution. The results are similar for Simulation 2 (Figure S17C&D): at low noise-levels (test-retest NSE < 0.4) noise correction corrects the bias introduced by noise while only modestly increasing the standard deviation. We limited our analyses to voxels with a test-retest NSE of less than 0.4 (measured using natural sounds in Paradigm I and both natural and model-matched sounds in Paradigm II), thus remaining in the regime where noise correction is well-behaved.

To directly test whether a test-retest NSE less than 0.4 is sufficient to ensure reliable measures, we measured the consistency of our noise-corrected measures across different subsets of data. Noise-correction requires two independent splits of data, and thus to test the reliability of noise-corrected NSE measures one needs at least 4 repetitions of each sound set. For Paradigm II, each subject heard between 6 and 15 repetitions of each sound set, which made it possible to perform this analysis. We averaged responses within four separate splits of data, each with an equal number of repetitions (e.g. assuming 12 repetitions, split 1 included repetitions 1, 5, 9, split 2 included repetitions 2, 6, 10 and so on). We then calculated the noise-corrected NSE twice based on splits 1 and 2 and splits 3 and 4. We excluded voxels with a test-retest NSE above 0.4 in splits 1 and 2 (because the test-retest NSE was only determined using splits 1 and 2, splits 3 and 4 provide a fully independent validation of the corrected values). This analysis revealed that the noise-corrected measures were reliable (Figure S18).

### Noise-correcting response variation and correlation measures

In addition to the normalized squared error, we also compared responses (in the six response components as well as in individual voxels) to natural and model-matched sounds by comparing their response variation, as measured by the standard deviation, and by correlating their responses (Figure 6E&F and Figure S12). We noise-corrected these measures as well. The variance of a noise-corrupted signal is biased upwards by the noise in the same manner as the signal power (equation 19), and thus can be corrected by subtracting off half of the residual power (the noise-corrected standard deviation can be computed by taking the square root of the noise-corrected variance). The correlation coefficient is given by:

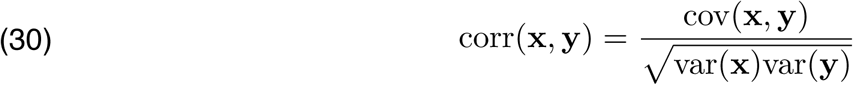

The covariance, which is defined as the cross-product of demeaned variables, is unbiased by noise for the same reason that the raw signal cross-product is unbiased by noise (equation 15), and thus we only need to correct the signal variance by subtracting off half of the residual power. This approach is similar to the more standard correction procedure of dividing by the square root of the test-retest correlation of the measures^45,85^ (and in the limit of infinite data the two are equivalent). However our approach is applicable when the test-retest reliability can only be measured for a single variable (as was the case for Paradigm I). MATLAB code implementing these noise-correction procedures can be downloaded here:https://github.com/snormanhaignere/general-analysis-code see noise_corrected_similarity.m

We note that the correlation between two variables becomes unstable (and in the limit undefined) as the variance of one variable approaches zero, which poses a problem in non-primary regions where we observed weak responses to the model-matched sounds. Thus, it was necessary to exclude voxels that did not have test-retest correlation to model-matched sounds of at least 0.4, which caused many non-primary voxels to be excluded in the maps of Figure S12. This is not an issue with the NSE, because the NSE is well-defined as long as either of the two variables being compared have non-zero variance.

### Model predictions with natural sounds

Our model assumes that voxels are a weighted sum of time-averaged statistics of the feature responses (equations 1-2). To predict voxel responses, we must choose a specific set of statistics and voxel weights. For the cochlear model, we used the average magnitude of each filter response’s envelope across time as our statistic (yielding 217 features, one per cochlear channel). For the three modulation models (temporal, spectral and spectrotemporal), we used the standard deviation of each feature’s response across time as our statistic, which we found gave better predictions than the power (sum of squares) or variance (sum of squares after demeaning) (we suspect this is because squaring the filters leads to a skewed distribution of values which are harder to linearly align with the voxel responses). We also included the 217 cochlear features in the modulation representation to make the analysis parallel to the model-matching procedure (where all of the modulation-matched sounds were also matched in their cochlear statistics). For the temporal modulation model, there were a total of 2170 features (9 rates × 217 audio frequencies + 217 cochlear channels). For the spectral modulation model there were 1736 features (7 scales × 217 frequencies + 217 cochlear channels). For the spectrotemporal modulation model, there were 27559 features (9 rates × 7 scales × 2 orientations × 217 frequencies + 217 cochlear channels). We did not include the temporal-only and spectral-only modulation features in the spectrotemporal modulation model because we found this did not improve prediction accuracy. We also excluded the DC filter from the temporal modulation model because it has zero variance across time.

For all of the models tested, we learned the voxel-specific weights across features via ridge regression, as is standard in the evaluation of encoding models^7,16^. Several of the models tested had a large number of features, which could potentially make it difficult to map the model features to the voxel responses. One option would have been to choose a subset of features or reduce dimensionality with PCA before performing regression. However, we have found that such approaches lead to slightly worse predictions than using a large number of features and regularizing with ridge^41^. Prior to regression, all of the features were normalized (z-scored across sounds). For models with both cochlear and modulation features, we separately re-scaled the two feature sets so that they would have the same norm and thus contribute similarly to the analysis (otherwise the modulation features would dominate because there were many more features in the modulation representation).

We used cross-validation across sounds to avoid statistical bias in fitting the weights as well as to select the optimal regularization parameter. First, we split the response of each voxel to the 36 natural sounds into test and train data. The training data was used to fit the weights and select the regularization parameter (details below), and the test data was used to evaluate the predictions of the model. We used 4-fold cross-validation, splitting the data into four equally sized sets of 9 sounds. For each set, we used the remaining sounds to the fit the model (i.e. the 27 sounds from the other three sets), and we averaged the accuracy of the predictions across the 4 folds. We quantified the similarity of the measured and predicted responses using the noise-corrected NSE so that the results could be compared with the model-matching results (details of noise correction given below).

To select the regularization parameter, we split each training set (27 sounds) again into four approximately equally sized sets (7, 7, 7, and 6). For each set, we used the remaining sounds to fit the weights for a large range of regularization parameters (2^-100^ to 2^100^ with octave steps), and the left out sounds to evaluate the accuracy of the model as a function of the regularization parameter. We then selected the regularization parameter that led to the best generalization accuracy averaged across the four splits (again using the noise-corrected NSE). Finally, given the selected regularization parameter, we fit the weights using all of the training set.

We used the same procedure for the cross-prediction analyses, but instead of training and testing on natural sounds, we learned the voxel weights on the natural sound responses and tested them on model-matched sounds (and vice versa). MATLAB code implementing these regression analyses can be downloaded here:

https://github.com/snormanhaignere/general-analysis-code, see: regress_predictions_from_3way_crossval.m regress_predictions_from_3way_crossval_noisecorr.m

### Noise-correcting model predictions

In the context of model predictions, we want to estimate the ability of the model to predict voxel responses to left out stimuli in the absence of noise due to fMRI. Because the predictions are derived from noisy fMRI measurements it is necessary to correct for the reliability of both the data and predictions^41^. Each natural sound was presented twice in the experiment. For each repetition and each test fold, we measured the response of each voxel to the test sounds and computed a prediction from the model using the training sounds (as described above in *Model predictions*). This procedure yielded two samples of the voxel response and two samples of the predicted response for each of the four test folds. We used these two samples to compute the necessary statistics for the noise-corrected NSE (equations 16-18 and 21-22). We used our noise-corrected squared error metric to both quantify the accuracy of the predictions and to select the regularization parameter.

### Voxel decomposition

Previously, we found that voxel responses to a diverse set of 165 natural sounds could be approximated by a weighted sum of a six canonical response patterns (components)^45^:

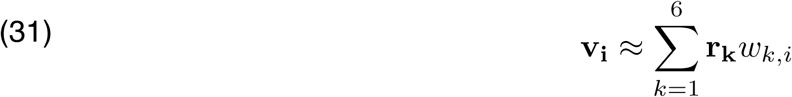

where **v_i_** and **r_k_** are 165-dimensional vectors representing the response of voxel *i* and component *k* to the sounds tested, and *w*_*k,i*_ represents the weight of component *k* in voxel *i*. The component responses (**r_k_**) and weights (*w*_*k,i*_) were jointly inferred by maximizing the non-Gaussianity of the weights, similar to classical independent component analysis^86^. Figure 6A re-plots a summary map of the weights from our prior study (averaged across subjects and transformed to a measure of statistical significance).

Six of the subjects from the present experiment also participated in our prior study, and two others participated in a similar experiment where we measured responses to a subset of 30 sounds from the original 165-sound experiment chosen to best identify the six components (by minimizing the variance of the component weights estimated by regression) (all eight subjects were scanned in Paradigm I). Four of these 8 subjects were scanned in the earlier version of the model-matching experiment without the spectral modulation condition, and thus the component responses to the spectral-only modulation-matched sounds were measured in just these four subjects. For each subject, we learned a set of reconstruction weights (*u*_*k,i*_) that when applied to the voxel responses from these two prior studies could approximate the component response profiles:

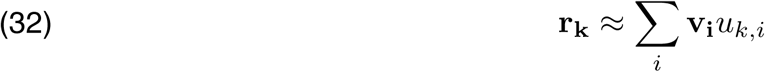

We then simply multiplied the voxel responses from the current experiment by the same reconstruction weights to estimate the component responses to the natural and model-matched stimuli from our current study. The reconstruction weights were estimated using ridge regression, picking the regularization parameter that led to the best prediction accuracy for left-out sounds (using the same cross-validation procedure described in the previous section to select the weights and regularization parameter; we used 5-fold cross-validation here). All voxels with a temporal SNR greater than 30 were used (temporal SNR was defined as the mean of the voxel’s timecourse divided by its standard deviation; results were similar when the analysis was restricted to voxels from the superior temporal plane / gyrus). This analysis was performed separately for every subject, and the inferred component responses were then averaged across subjects (this made it possible to use bootstrapping to compute standard errors and significance, see *Statistics* below). We again quantified the similarity of responses to natural and model-matched sounds using the noise-corrected NSE.

We note that an alternative approach would have been to use the pseudoinverse of the encoding weights (*w*_*k,i*_ in equation 31) as our reconstruction weights^45^, rather than learning reconstruction weights via ridge regression. We have consistently found the pseudoinverse approach to be less effective than directly learning the reconstruction weights (i.e. the reconstructed profiles more closely match the target response profile in left-out data when the reconstruction weights are directly optimized for the purpose of reconstruction). The approach of learning separate weights for the purpose of reconstruction is standard in the sensory encoding/decoding literature^87^.

### Annular analyses

We quantified the similarity of responses to natural and model-matched sounds (or model predictions) by binning voxels based on their distance to PAC, defined either tonotopically or anatomically. Voxels were binned in 5 mm intervals, and we computed the median NSE value across the voxels within each bin. Anatomically, we defined PAC as the center of TE1.1, which is located in posteromedial Heschl’s Gyrus. We relied on surface-based alignment to map the TE1.1 ROI to the appropriate anatomical region (the presence/absence of duplications along HG is reported in Table S1; defined by inspection using the scheme described in Da Costa et al. (2011)^52^). Tonotopically, PAC was defined by hand in individual subjects as the center of the low-frequency reversal of the high-low-high gradient within Heschl’s Gyrus^51–55^. These maps were derived from responses to pure tones presented in six different frequency ranges (with center frequencies of 200, 400, 800, 1600, 3200, & 6400). We measured the frequency range that produced the maximum response in voxels significantly modulated by frequency (p < 0.05 in a 1-way ANOVA across the 6 ranges); the details of the stimuli and analyses have been described previously^81^. Group tonotopy maps were based on a cohort of 21 subjects who were run in this tonotopy localizer across multiple studies (6 of the subjects from this experiment were part of this cohort)^88^. The best-frequency maps from each of these 21 subjects were averaged to form group maps. Voxels in which fewer than three subjects had frequency-modulated voxels were excluded from the map.

### Statistics

All of our statistical tests with the exception of the voxel decomposition analysis (Figure 6) were based on the annular analyses described above. We defined primary and non-primary regions using the three bins nearest and furthest from PAC, defined anatomically or tonotopically. We then used standard random effects analyses (t-tests) to compare the NSE values of primary and non-primary regions. In every subject and hemisphere we observed an increase in the NSE between primary and non-primary regions, which was significant using a t-test as well as a simple sign test. The same was true for comparing NSE values derived from model matching and model prediction: in all eight subjects, the increase in NSE values between primary and non-primary regions was greater for model-matching than for prediction.

For comparing NSE values between different model-matching conditions, we were only able to compute individual-subject maps from the four subjects that were scanned multiple times in Paradigm I. All four subjects tested showed the trends evident in the group map (Figure S7), but the small number of subjects precluded a random effects analysis. We thus performed statistics on group-averaged responses, bootstrapping across all 12 subjects scanned at least once in Paradigm I. Specifically, we sampled 12 subjects with replacement 1000 times, and averaged responses across the 12 sampled subjects in standardized anatomical coordinates. For each sample, we then recomputed the voxel-wise NSE values and binned these values based on distance to PAC. This procedure yielded 1000 samples of the NSE for each annular bin and condition. For each contrast (e.g. NSE full model < NSE cochlear matched sounds), we subtracted the average NSE across all bins between the two conditions being compared, and counted the fraction of times this contrast fell below zero (yielding the reported p-values). Results were similar when we averaged responses within just PAC (the first three bins) where the model performed best.

An obvious downside of statistics based on group-averaged maps is that individual subjects exhibit idiosyncrasies in their anatomy^89^. Component analysis provides one way to overcome this problem by mapping all of the subjects to a common response space, an approach that is relatively common in EEG studies^90^ but is less frequently applied to fMRI analyses^91^. We performed stats on the component responses by estimating the response of each component in each subject, and then bootstrapping across the eight subjects in which we had component data. We randomly sampled 8 subjects with replacement 1000 times, averaged the component responses for those subjects, and recomputed the noise-corrected NSE. We then contrasted NSE values between components or conditions to assess significance.

---

**Figure S1.**
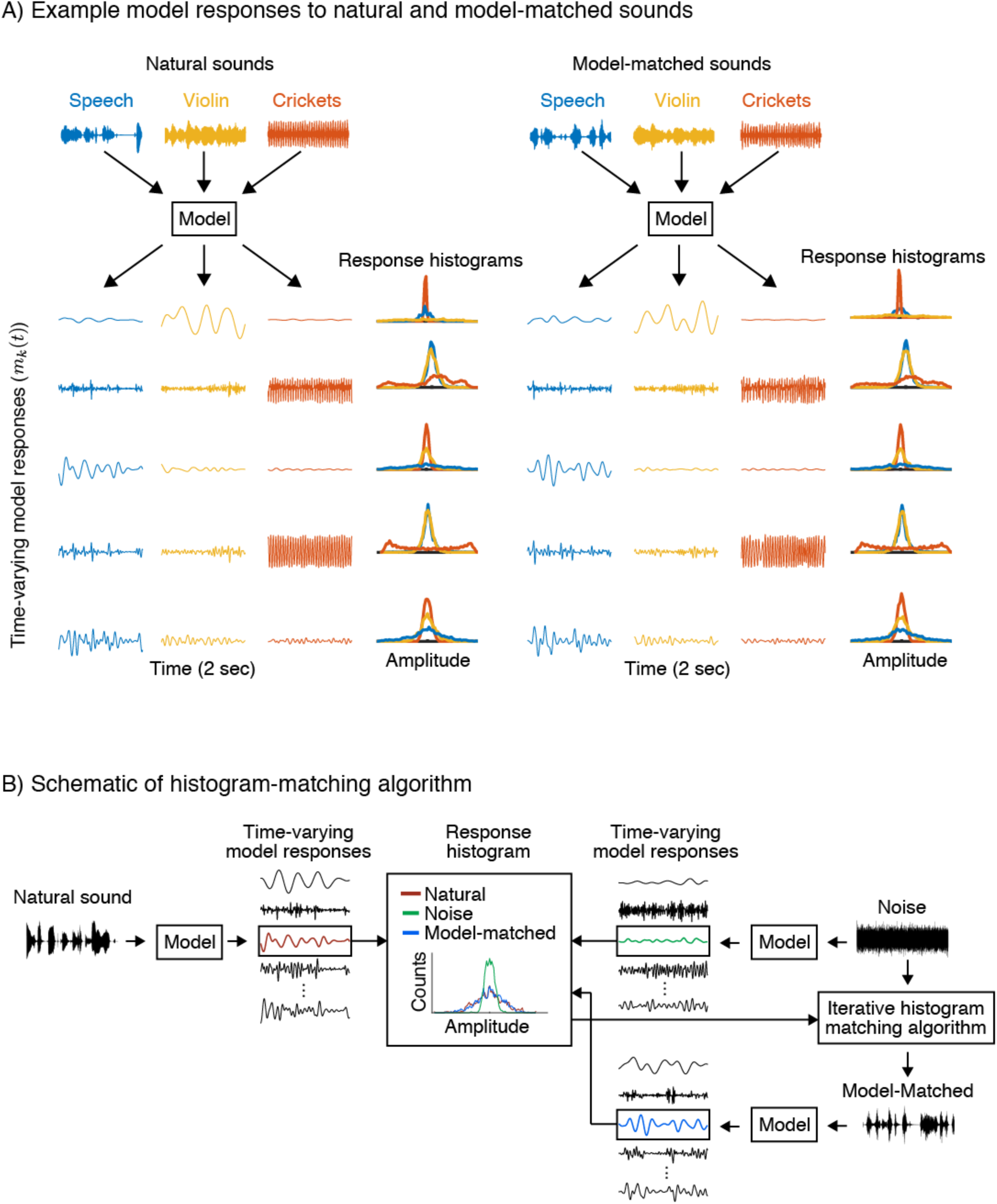
Schematic of model-matching approach. (A) All of the models considered are defined by the response timecourse of a set of model features, each computed by filtering a cochleagram representation of sound (illustrated in Figure 1A). Our model-matching algorithm collapses these timecourses across time to form a histogram, and then generates a sound with the same histograms as a natural sound. Here we plot example timecourses and histograms for three example natural sounds (left panel) and corresponding model-matched sounds (right panel). Different natural sounds produce distinct response timecourses and histograms. Corresponding natural and model-matched sounds produce similar response histograms, but distinct response timecourses. (B) The model-matched sounds were synthesized by modifying a noise signal so as to match the histogram of each feature’s response to a natural sound. The algorithm was initialized with Gaussian noise that was initially unstructured, and thus produced feature responses with a different histogram and a different time-varying response pattern. The noise sound was then iteratively adjusted so as to match the histogram of each feature to the natural sound, while leaving the temporal pattern unconstrained. This figure plots histograms for one example model feature in response to a natural, noise and model-matched sound. The histogram matching algorithm is conceptually similar to a classic visual texture synthesis algorithm^48^.

**Figure S2.**
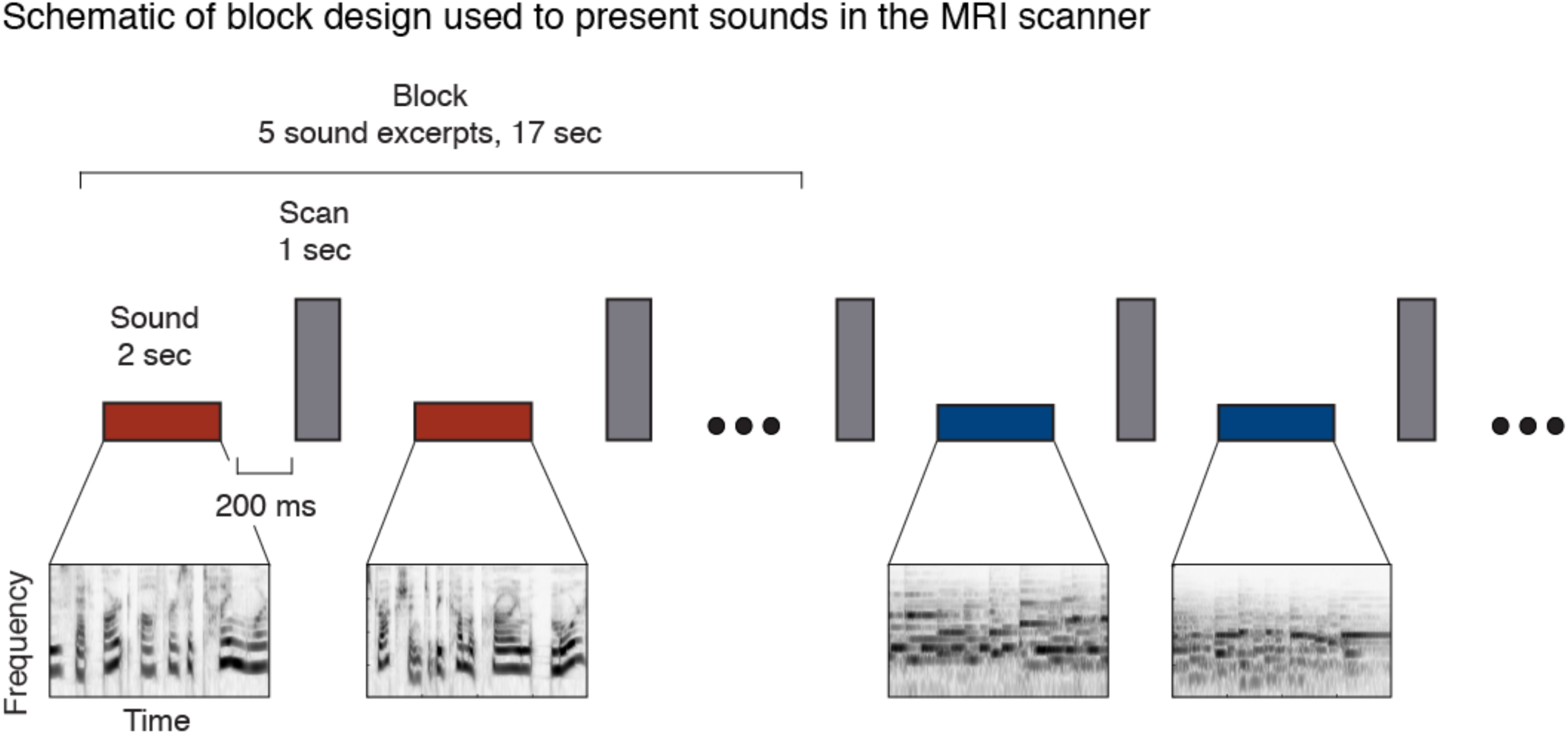
Schematic of the sparse scanning paradigm used to present stimuli in the experiment. Each 10-second stimulus was subdivided into five 2-second segments. These five segments were presented in a random order with a 1-second scan acquisition interspersed between each presentation (1.05 seconds for Paradigm II). A short 200-ms buffer was present between stimuli and scan acquisitions. The total duration of each “block” of 5 sounds was 17 seconds (17.25 seconds for Paradigm II). The response to a stimulus was computed as the average response of the 2^nd^ through 5^th^ scan acquisition after block onset (the first acquisition was discarded to account for the hemodynamic lag).

**Figure S3.**
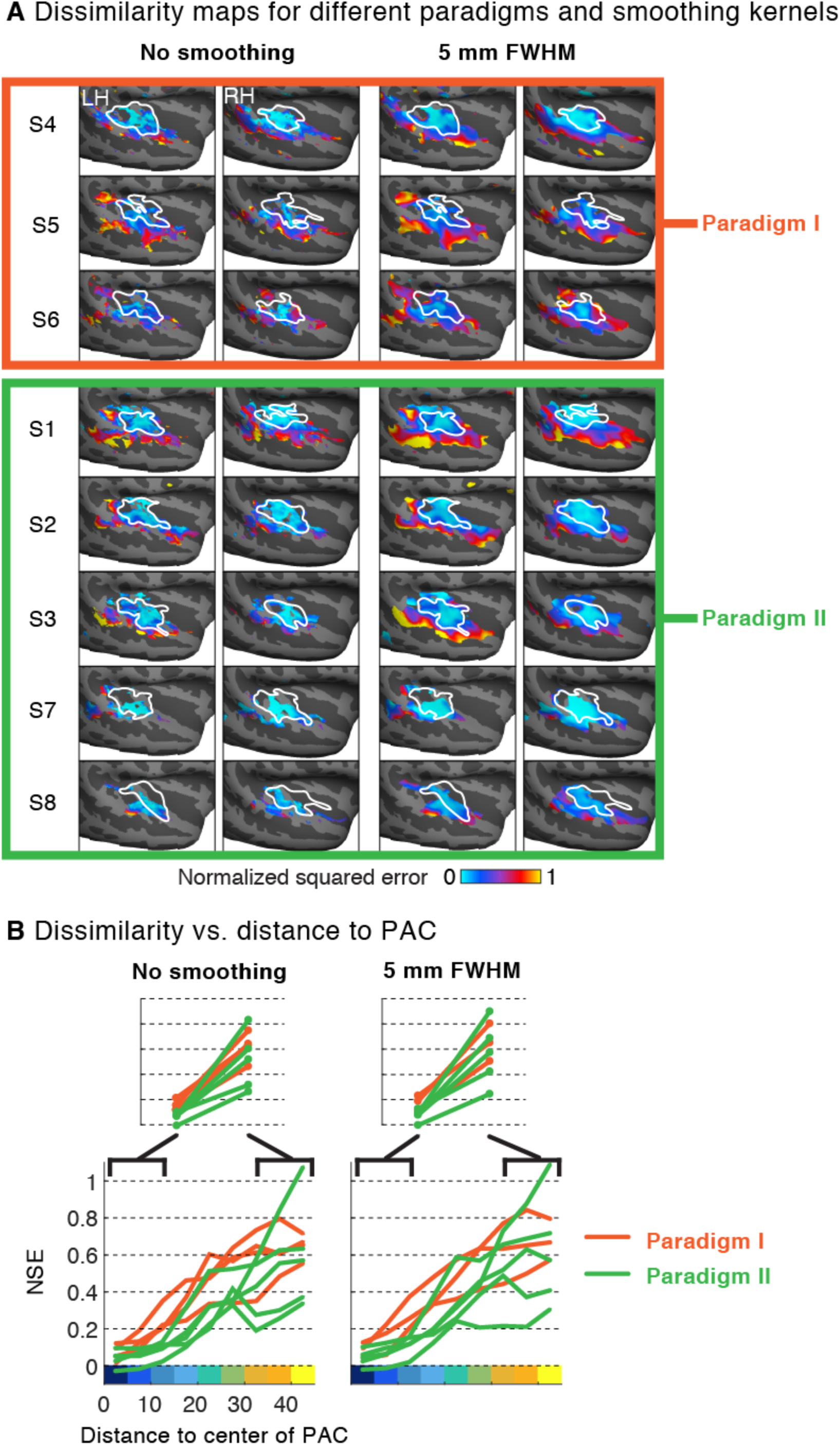
Results broken down by paradigm and the presence/absence of smoothing. In Paradigm I, only the natural sounds were repeated. In Paradigm II, both natural and model-matched sounds were repeated. A smaller voxel size was employed in Paradigm II (2 mm isotropic instead of 2.1 × 2.1 × 4 mm for Paradigm I). (A) Natural vs. model-matched dissimilarity maps computed with and without smoothing. Individual subjects are grouped by Paradigm. Subjects are sorted by the reliability of their response to natural sounds for Paradigm I and by the reliability of their response to both natural and model-matched sounds for Paradigm II (measured using the NSE). (B) Annular analyses computed from data with and without smoothing. Each line corresponds to an individual subject and the color indicates the paradigm (orange for Paradigm I and green for Paradigm II). NSE values are averaged across the left and right hemisphere since we observed similar trends in both hemispheres.

**Figure S4.**
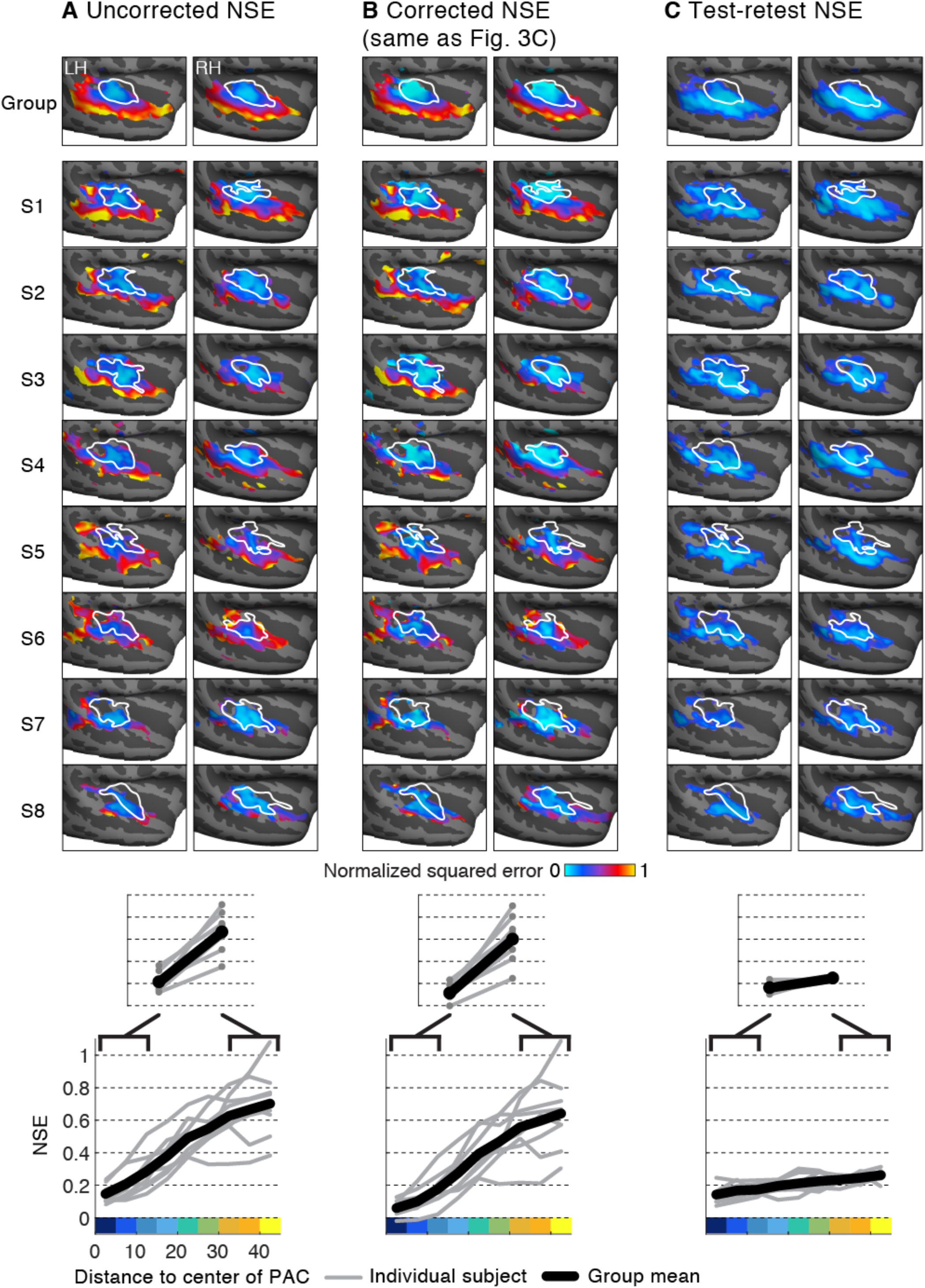
Noise correction and voxel reliability. (A) The uncorrected NSE between responses to natural and model-matched sounds. (B) Corrected NSE maps (same as Figures 3C) replicated here for ease of comparison with the uncorrected maps. Test-retest reliability of voxel responses measured with the NSE. Voxel reliability for Paradigm I (Group, S4, S5, S6) is based on responses to natural sounds. Voxel reliability for Paradigm II (S1, S2, S3, S7, S8) is based on responses to both natural and model-matched sounds (responses to natural and model-matched sounds were combined into a single vector, and we computed the NSE for multiple measurements of this vector). Distance-to-PAC analyses are shown at the bottom of each panel. Format is the same as Figure 3C&D.

**Figure S5.**
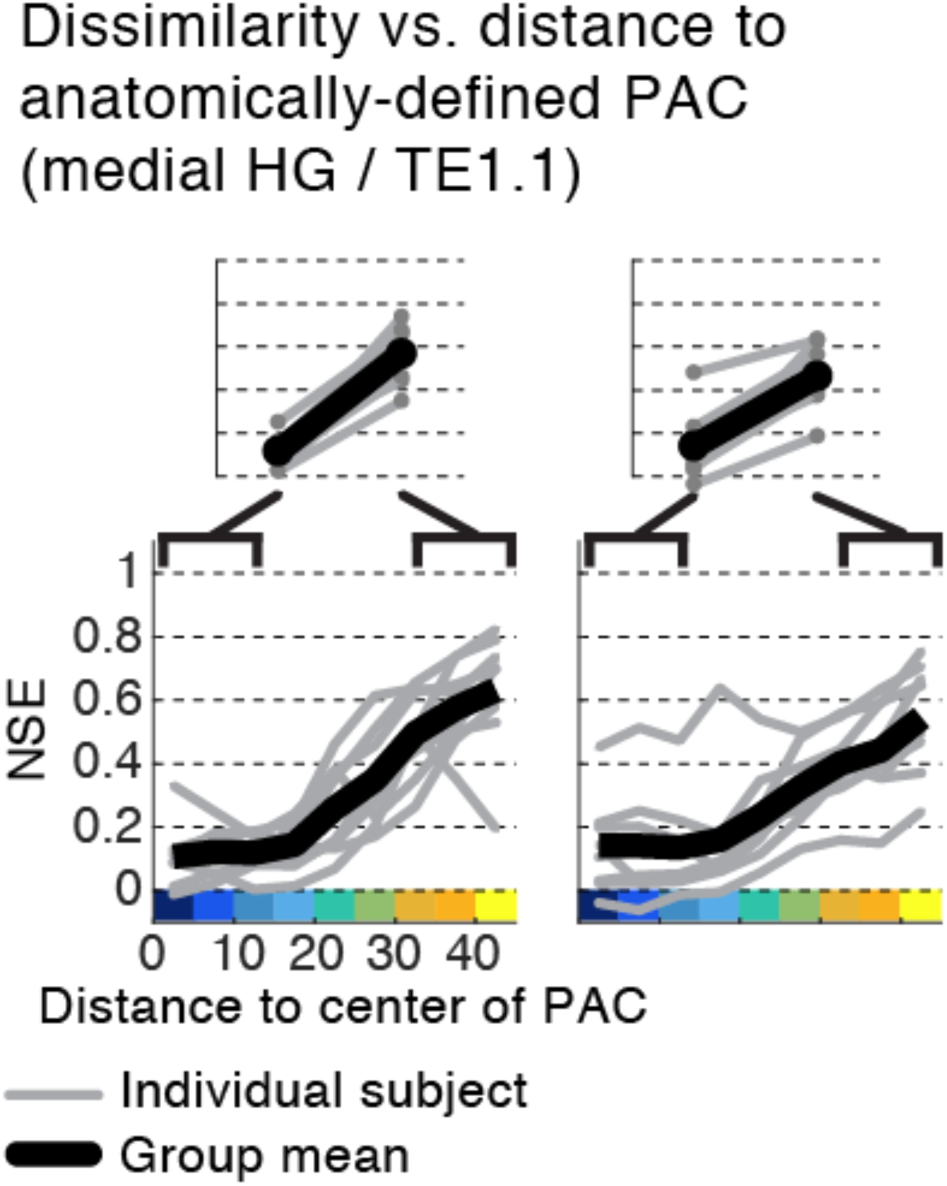
Dissimilarity vs. distance from PAC using an anatomical rather than tonotopic definition of PAC. Voxels are binned based on their distance to center of anatomical region TE1.1^58^, which is located in posteromedial HG. Format is the same as Figure 3D.

**Figure S6.**
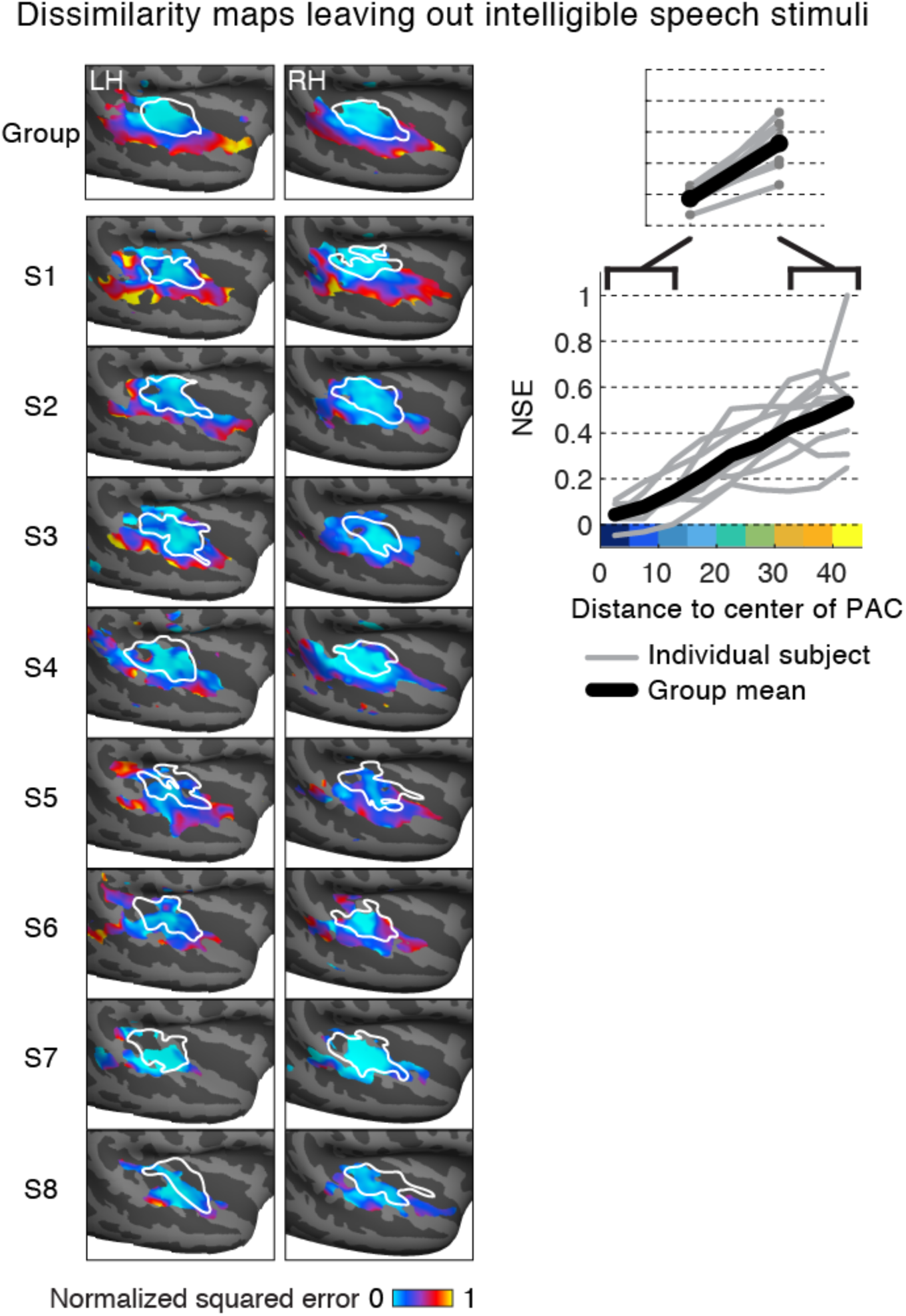
Dissimilarity maps and annular analyses omitting intelligible speech stimuli. Maps plot the normalized squared error (NSE) between voxel responses to natural and model-matched sounds omitting English speech and music with English vocals (all subjects were native English speakers). Format is the same as Figure 3C&D.

**Figure S7.**
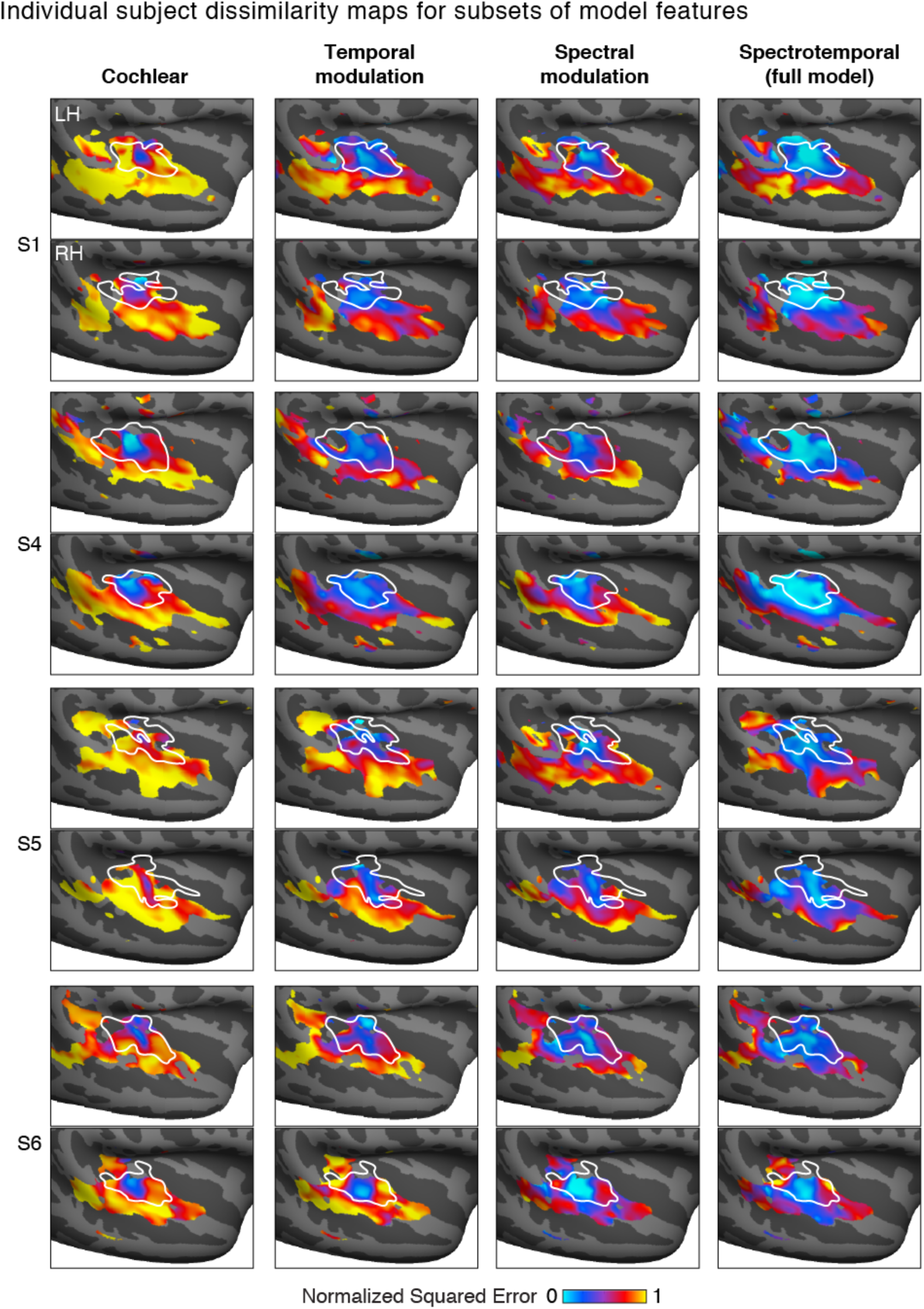
Individual subject maps of dissimilarity between responses to natural and model-matched sounds (NSE) for subsets of model features. Format the same as Figure 4B. Only subjects scanned in Paradigm I are shown, since Paradigm II did not include model-matched sounds constrained by subsets of model features.

**Figure S8.**
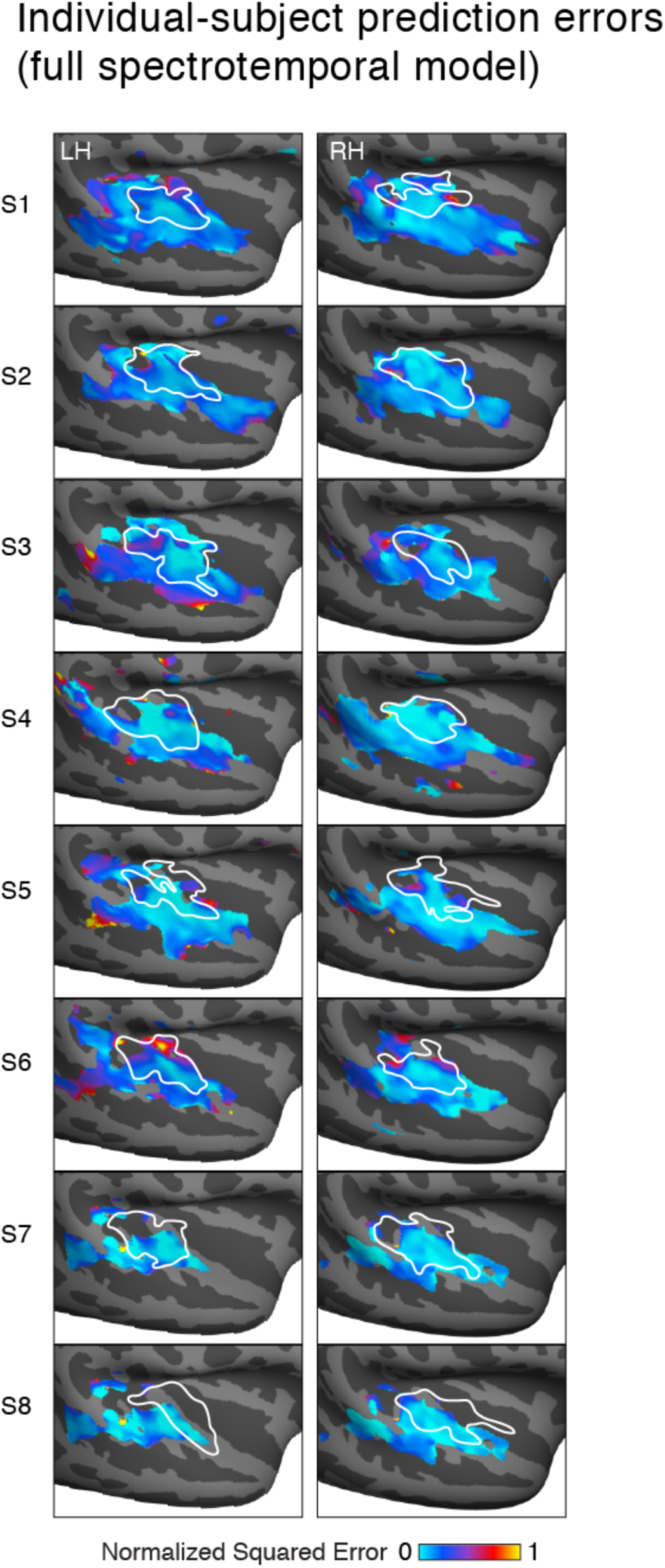
Individual subject prediction error maps based on the full spectrotemporal modulation model (maps for subsets of model features are omitted due to space constraints, but, like the maps for the full model, they resembled those of the group). Format the same as Figure 3C.

**Figure S9.**
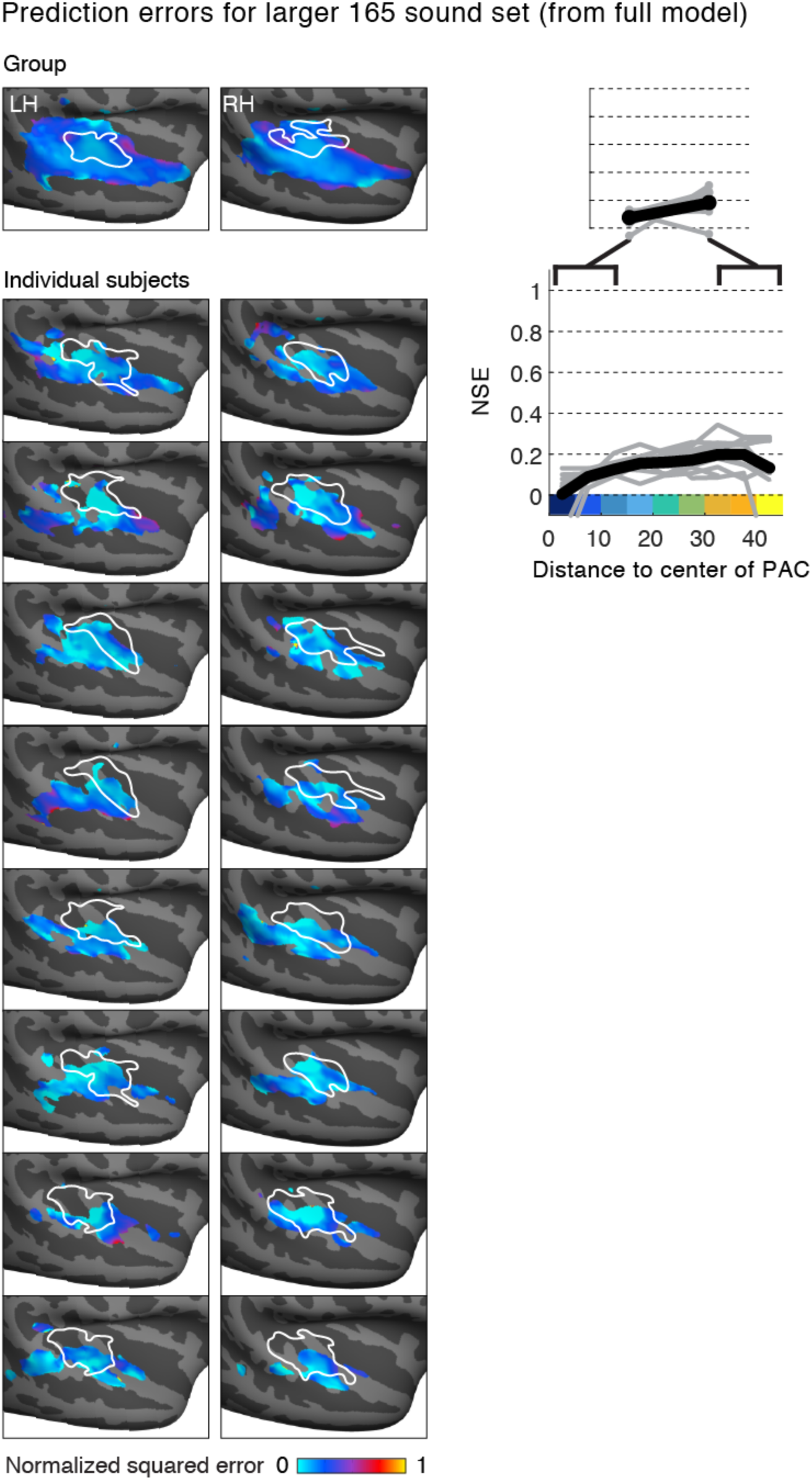
Prediction accuracy of the full spectrotemporal model using a larger set of 165 natural sounds tested in a prior study. Format is the same as Figure 4B&C.

**Figure S10.**
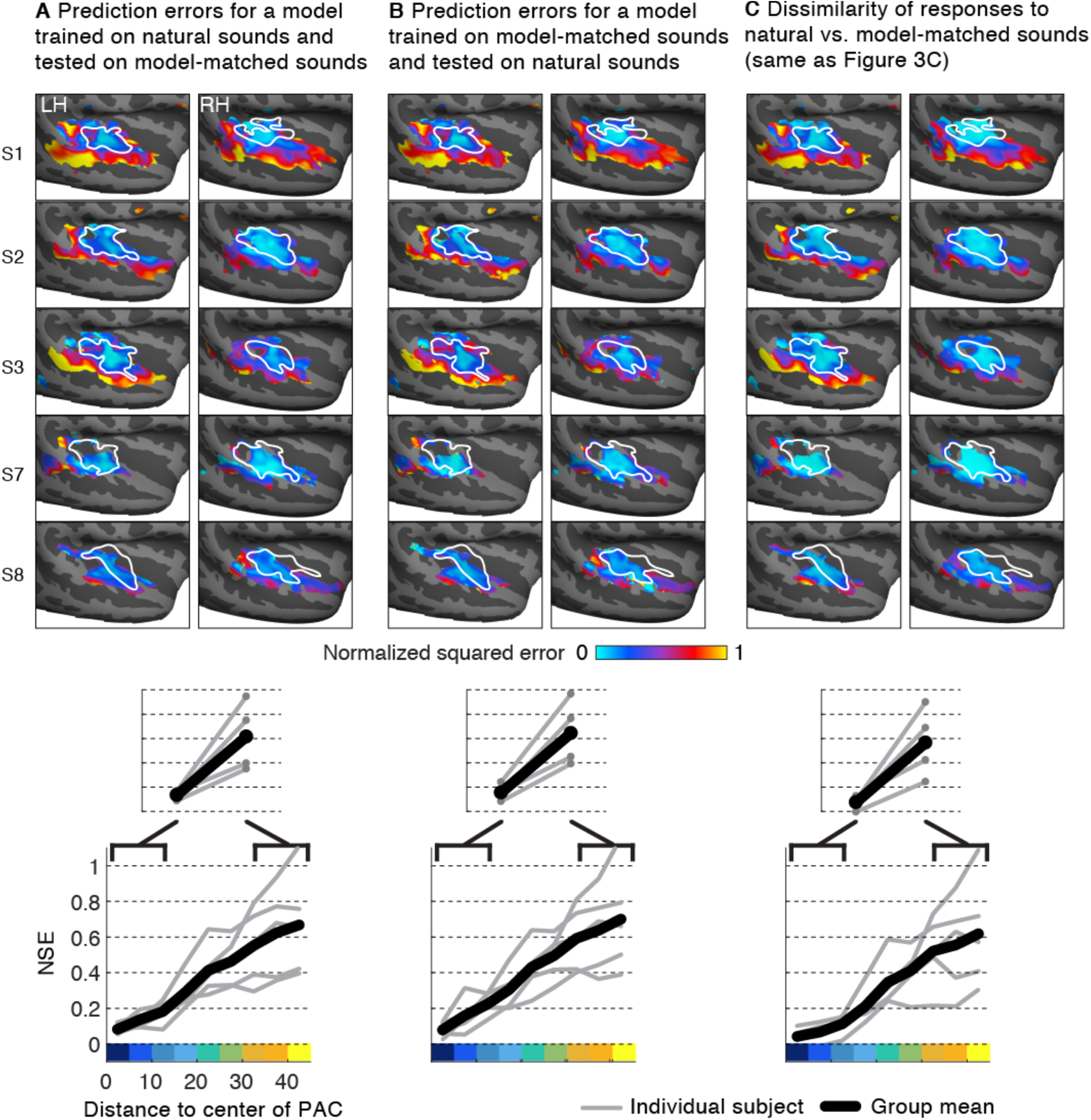
(A) Prediction error maps of a model trained on natural sounds and tested on model-matched sounds. (B) Prediction error maps of a model trained on model-matched sounds and tested on natural sounds. (C) For comparison, the error of the measured voxel response to natural and model-matched sounds is reproduced here (same as Figure 3C&D). Annular analyses summarizing the error as a function of distance to tonotopically-defined PAC are shown below each set of maps. Data are shown for subjects scanned in Paradigm II for which both natural and model-matched sounds were repeated, which made it possible to noise-correct the predictions.

**Figure S11.**
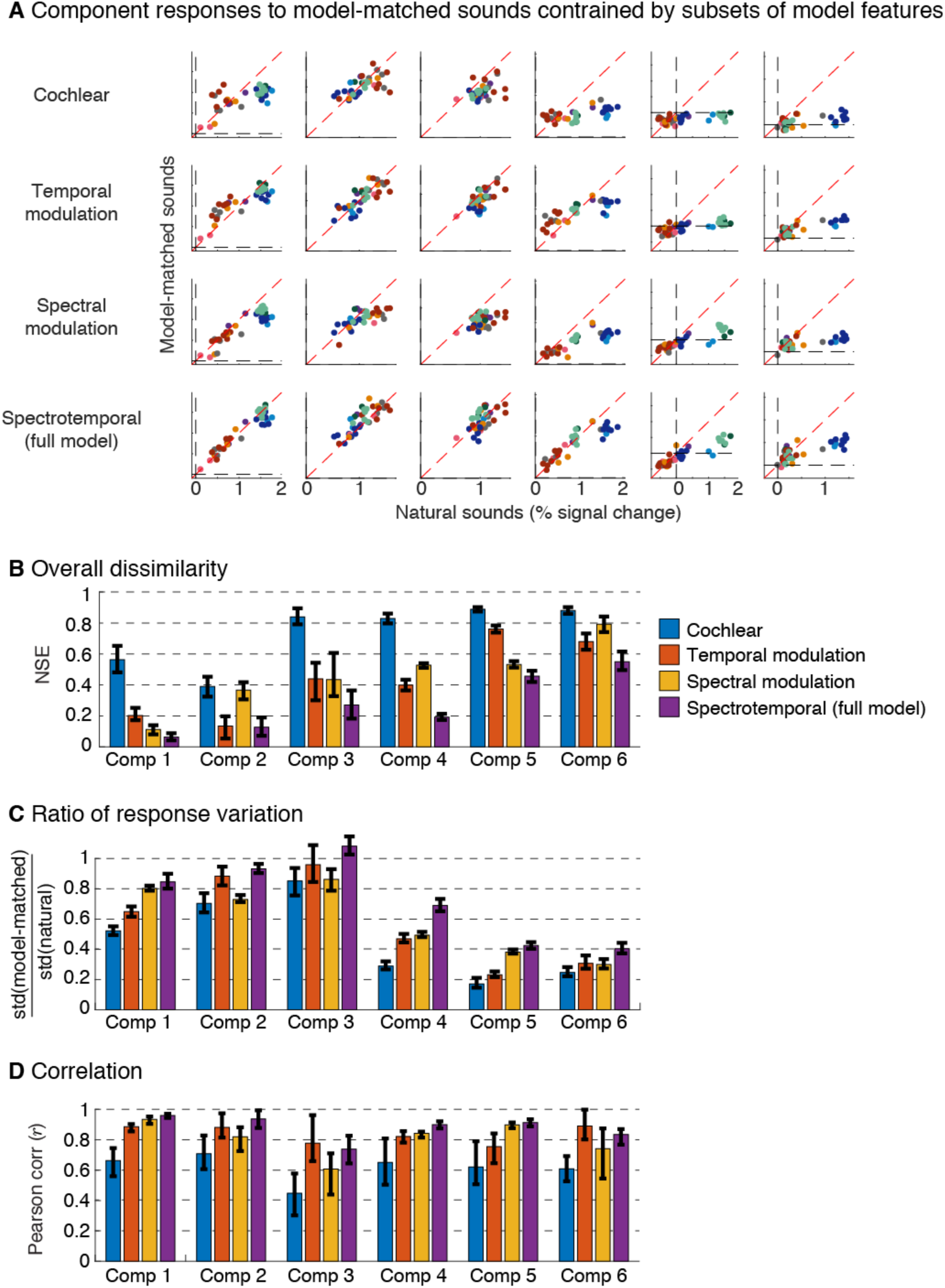
Plots showing component responses to sounds constrained by subsets of model features, as well as the full model (Figure 6 only shows results from the full model). (B) Normalized squared error between natural and model-matched sounds. (C) Ratio of the standard deviation of responses to model-matched and natural sounds. (D) Correlation of responses to natural and model-matched sounds. Format is the same as Figures 6C-6F.

**Figure S12.**
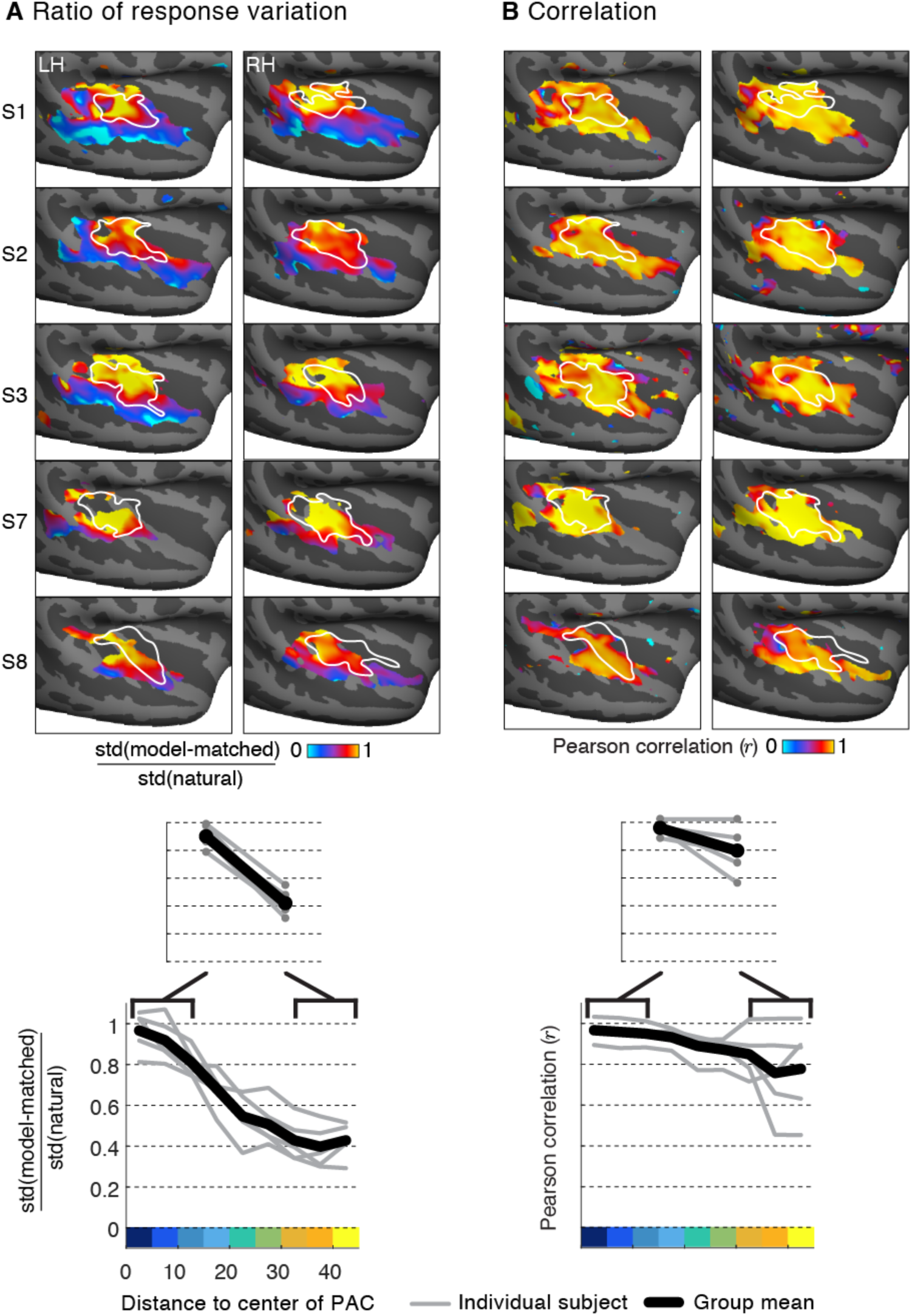
Examination of the nature of the divergent responses to natural and model-matched sounds. (A) Whole-brain maps plotting the variation in responses to natural vs. model-matched sounds, measured as the ratio of the standard deviation of responses to the two sound sets. Cool colors indicated less response variation for model-matched sounds. Distance-to-PAC summary analysis is plotted below. (B) Maps of the Pearson correlation between responses to natural and model-matched sounds with distance-to-PAC analysis below. All of the measures have been corrected for noise. Analysis is based on data from Paradigm II in which we measured responses to natural and model-matched sounds an equal number of times. For the response variation maps (panel A), we included all voxels with a reliable response across both natural and model matched sounds (NSE < 0.4). For the correlation maps (panel B), we excluded voxels that did not have a reliable correlation to model-matched sounds (r < 0.4), as was the case in many non-primary voxels due to weak responses. For such voxels, it is difficult to estimate a reliable correlation, since the correlation is undefined as the variance of one variable goes to zero.

**Figure S13.**
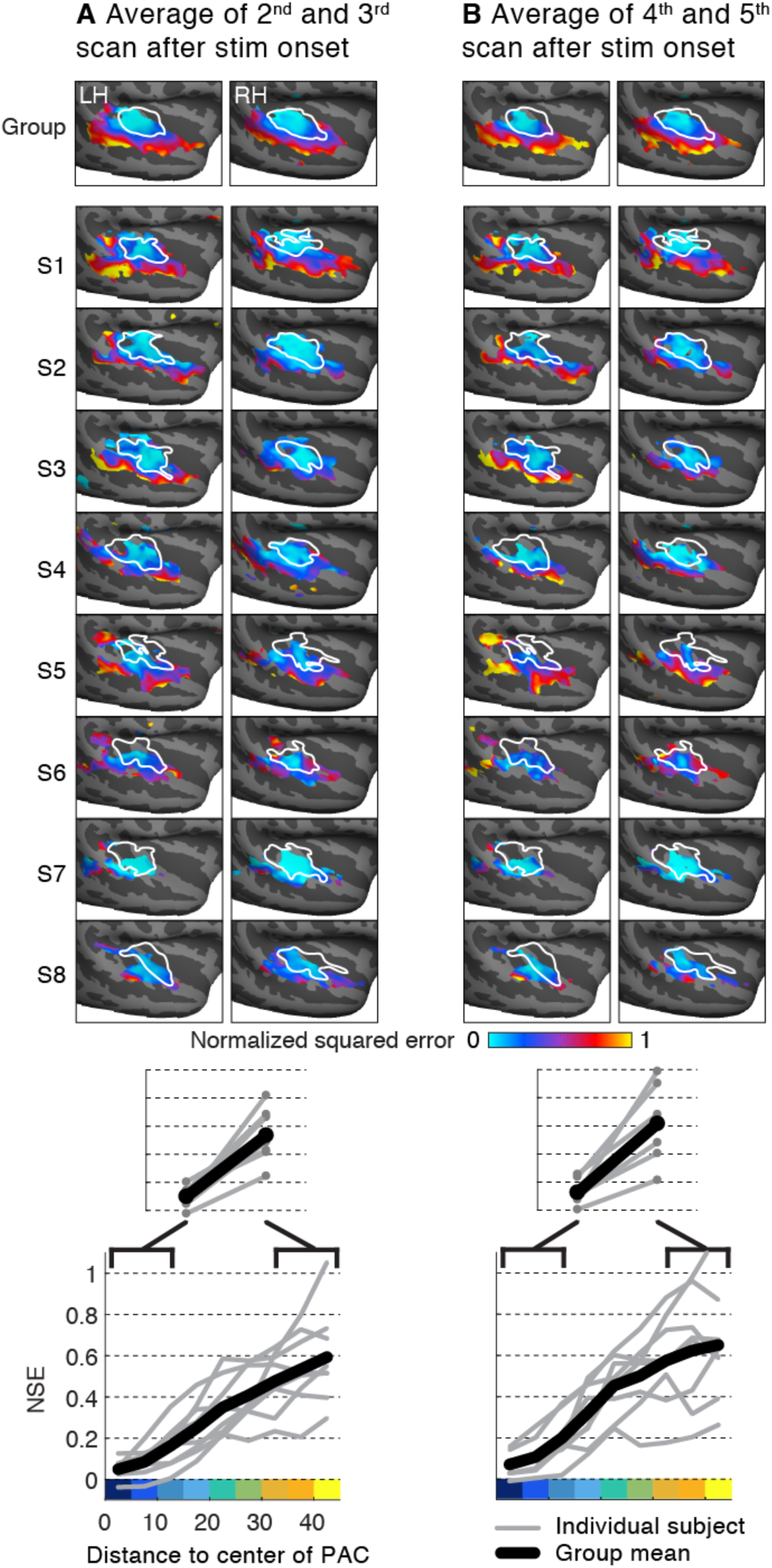
Effect of the fMRI averaging window on the dissimilarity of responses to natural and model matched sounds. Each stimulus was 10-seconds in duration, but was split-up into five 2-second segments (see Figure S2). After each segment a single scan acquisition was collected. Analyses in the main text were based on the average response of the 2^nd^ through 5^th^ acquisition after the onset of each stimulus block (first acquisition was discarded to account for the hemodynamic delay). Here we test the sensitivity of the results to the averaging window by restricting the analysis to data averaged across acquisitions 2 and 3 (panel A) or 4 and 5 (panel B). Compare with Figures 3C&D.

**Figure S14.**
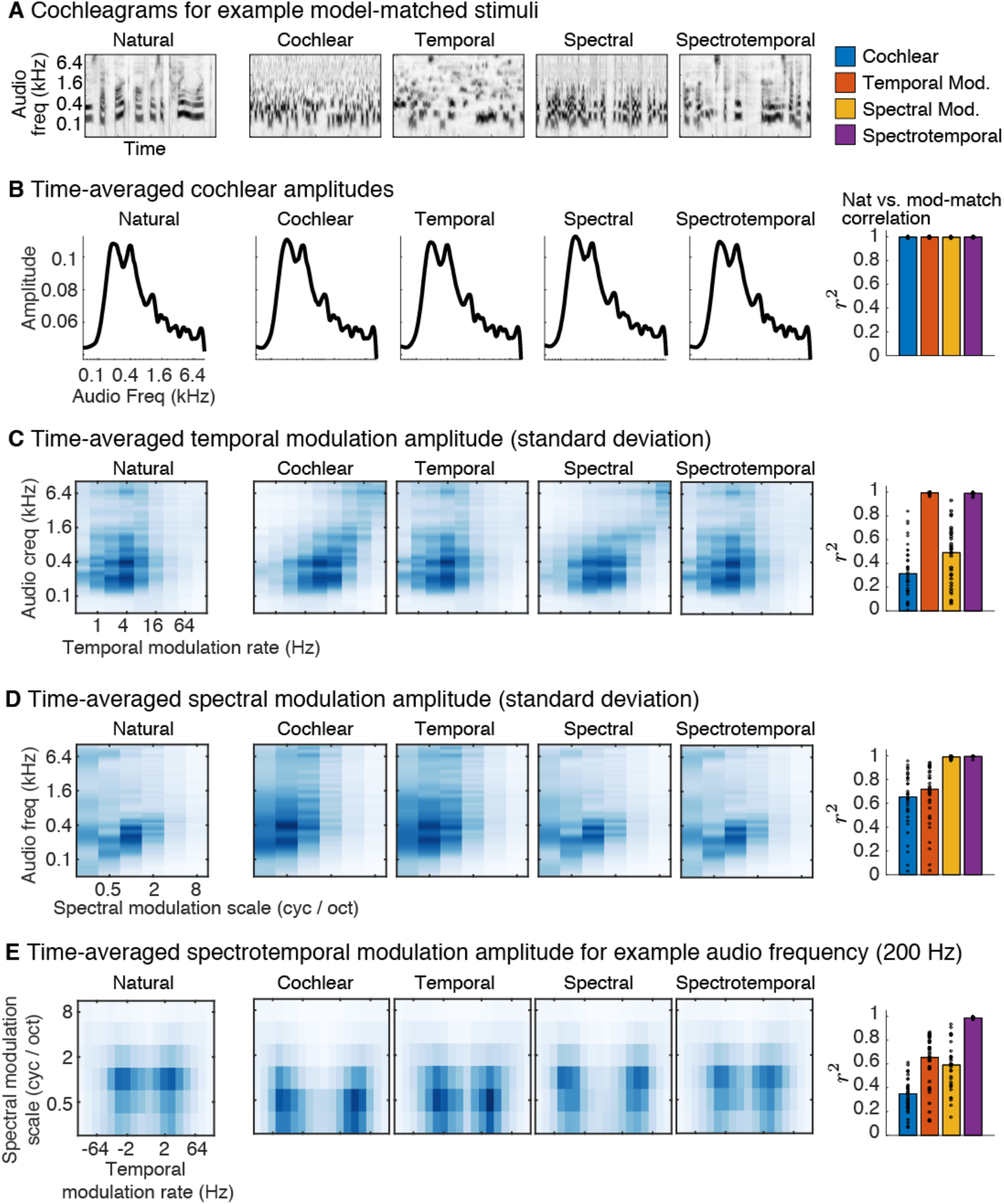
Validation of the model-matching synthesis procedure via comparison of time-averaged statistics for natural and model-matched sounds. (A) Cochleagrams for a natural sound (a speech excerpt) and four corresponding model-matched sounds. (B-E) Each model was defined by a set of feature responses. This figure plots a time-averaged measure of the amplitude of each feature’s response to the example natural and model-matched sounds shown in panel A. The right side of the panel plots the correlation of the filter amplitudes across all model filters for corresponding natural and model-matched sounds. Each point corresponds to a single pair of natural/model-matched sounds. (B) Amplitude of each cochlear frequency channel envelope averaged across time. Cochlear channel power is matched in all four conditions, as desired/expected. (C) Temporal modulation amplitude (standard deviation of each feature across time) for example natural and model-matched sounds. Modulation amplitude is plotted as a function of the filter’s preferred audio frequency and temporal modulation rate. Spectral modulation amplitude plotted as a function of the filter’s preferred audio frequency and spectral modulation scale. Spectrotemporal modulation amplitude plotted as a function of temporal modulation rate and spectral modulation scale for an example audio frequency (200 Hz).

**Figure S15.**
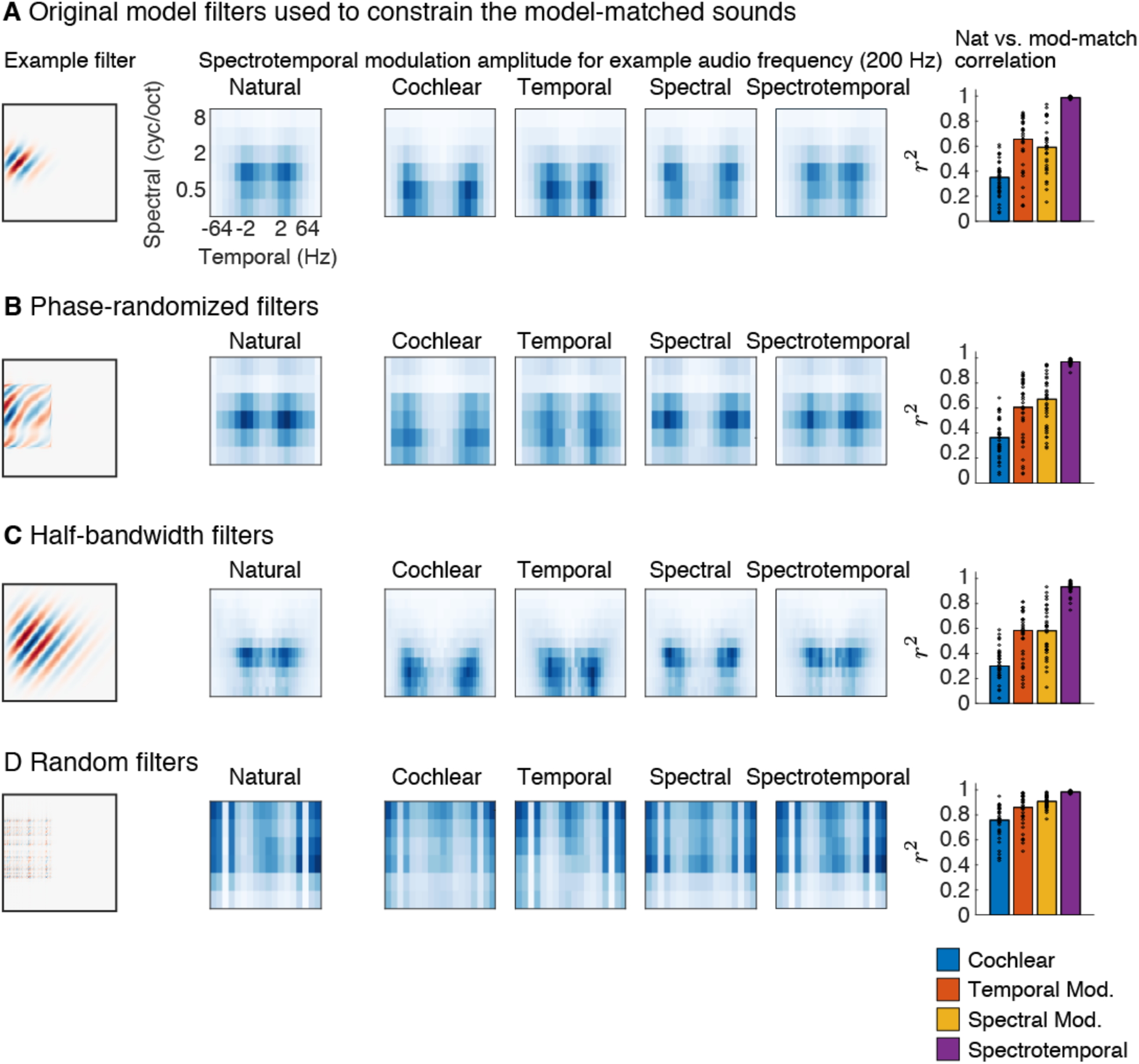
Comparison of how well the natural and synthesized (model-matched) sounds are matched when evaluated using spectrotemporal filters that differed from those used to generate the model-matched sounds. In each case, we plot an example filter from the model (left), the amplitude (standard deviation) of the filter responses as a function of the temporal rate and spectral scale for an example audio frequency (200 Hz) (middle), and the correlation of the amplitude across all of the filters for the natural and model-matched sounds (right) (format similar to Figure S14E). (A) The original spectrotemporal filters from Chi et al. (2005) that were used to constrain the model-matched sounds (same as Figure S14E). (B) Spectrotemporal filters with randomized temporal and spectral phase. (C) A model with narrower bandwidths and more filters to compensate. (D) A random filter basis with variable temporal and spectral extent. In all four cases, the measured modulation power is similar for the natural and model-matched sounds. This suggests that voxels with similar responses to natural and model-matched sounds are compatible with a wide range of spectrotemporal modulation filters, and that a wide range of such filters are ruled out as descriptions of voxels that give different responses to natural and model-matched sounds, like those we observed in non-primary regions.

**Figure S16.**
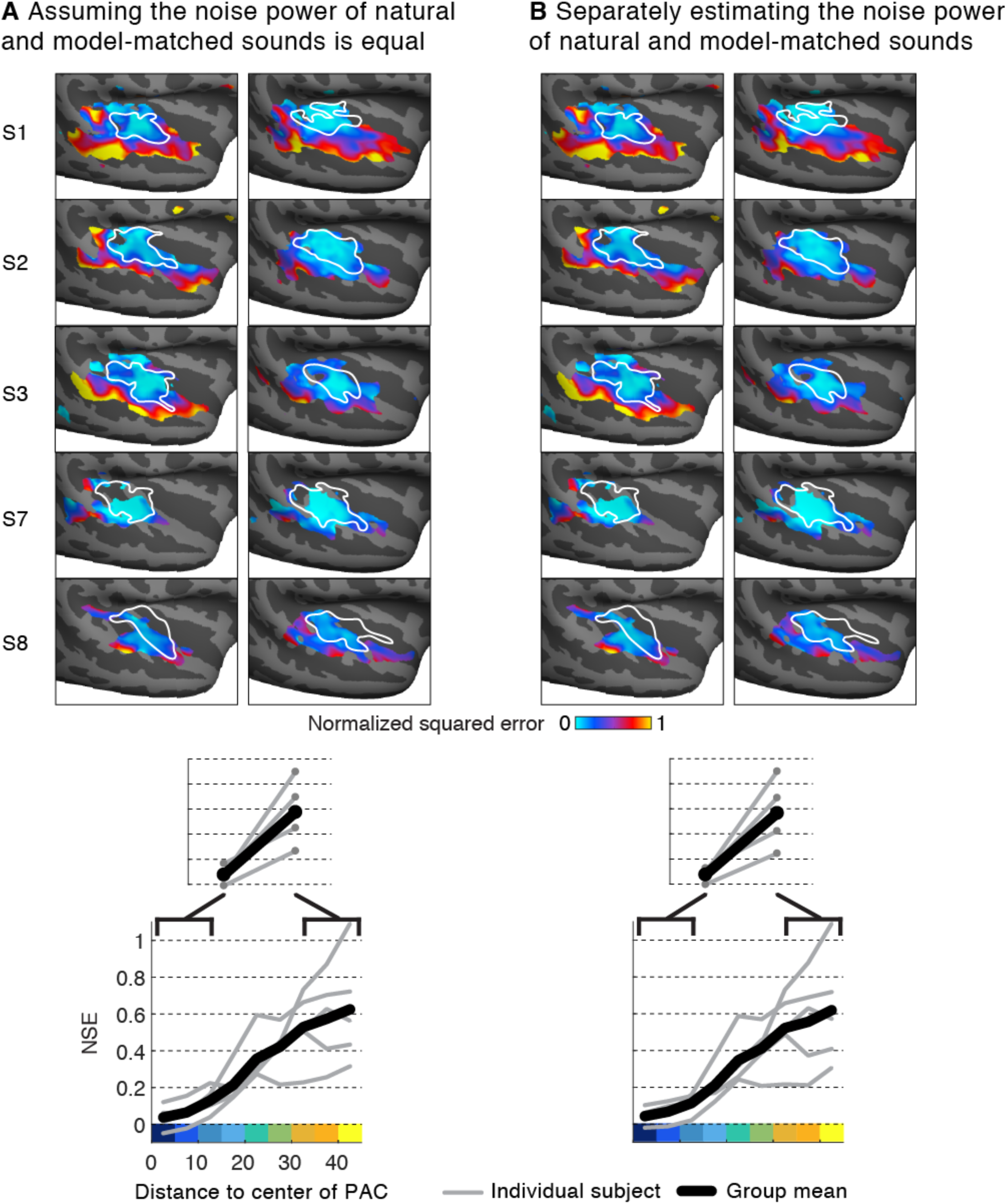
Comparison of noise correction using noise estimates exclusively from responses to natural sounds or from both natural and model-matched sounds. Noise-correction required estimating the power of the noise for natural and model-matched sounds. For Paradigm I, only responses to natural sounds were repeated in each scan. Using data from Paradigm II, we tested whether it is necessary to separately estimate the noise power for natural and model-matched sounds, or whether one can assume they are equal. (A) Noise-corrected NSE values computed by assuming the noise power for natural and model-matched sounds is equal, and using only responses to natural sounds to compute it. (B) Noise-corrected NSE values computed by separately estimating the noise power for natural and model-matched sounds (same maps as those in Figure 3C). Results are similar in both cases.

**Figure S17.**
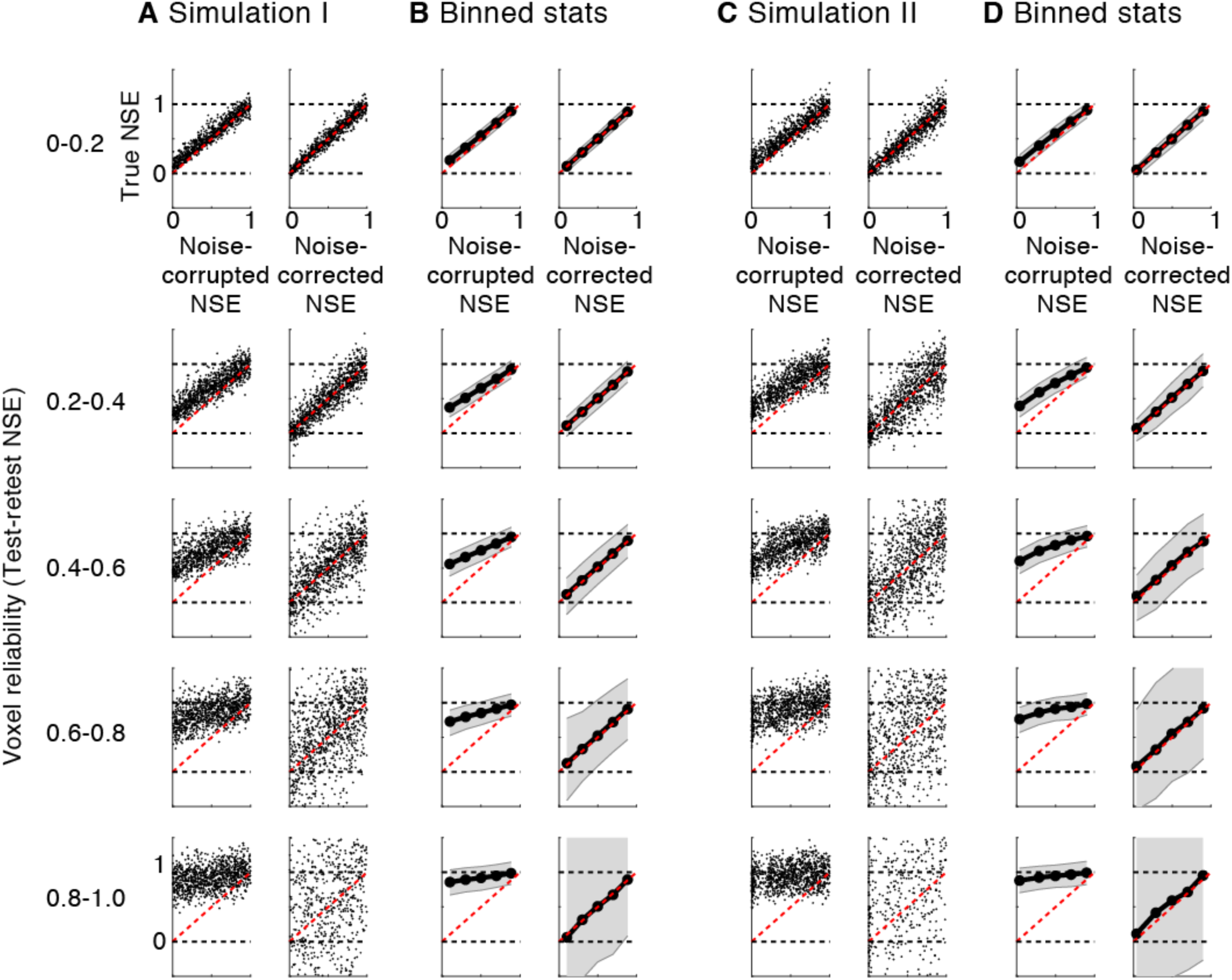
Results of noise-correction simulations. (A) Each dot corresponds to a single simulated voxel. The true NSE value is plotted against the noise-corrupted and noise-corrected NSE values. Results have been grouped by the reliability of the simulated voxel responses as measured by the test-retest NSE of the voxel responses (from high to low reliability going from top to bottom). (B) The median and standard deviation of the samples. (C&D) Same as for panels A&B, but for Simulation II (see *Evaluating the noise-corrected NSE with simulated data* in the Methods for details of the two simulations).

**Figure S18.**
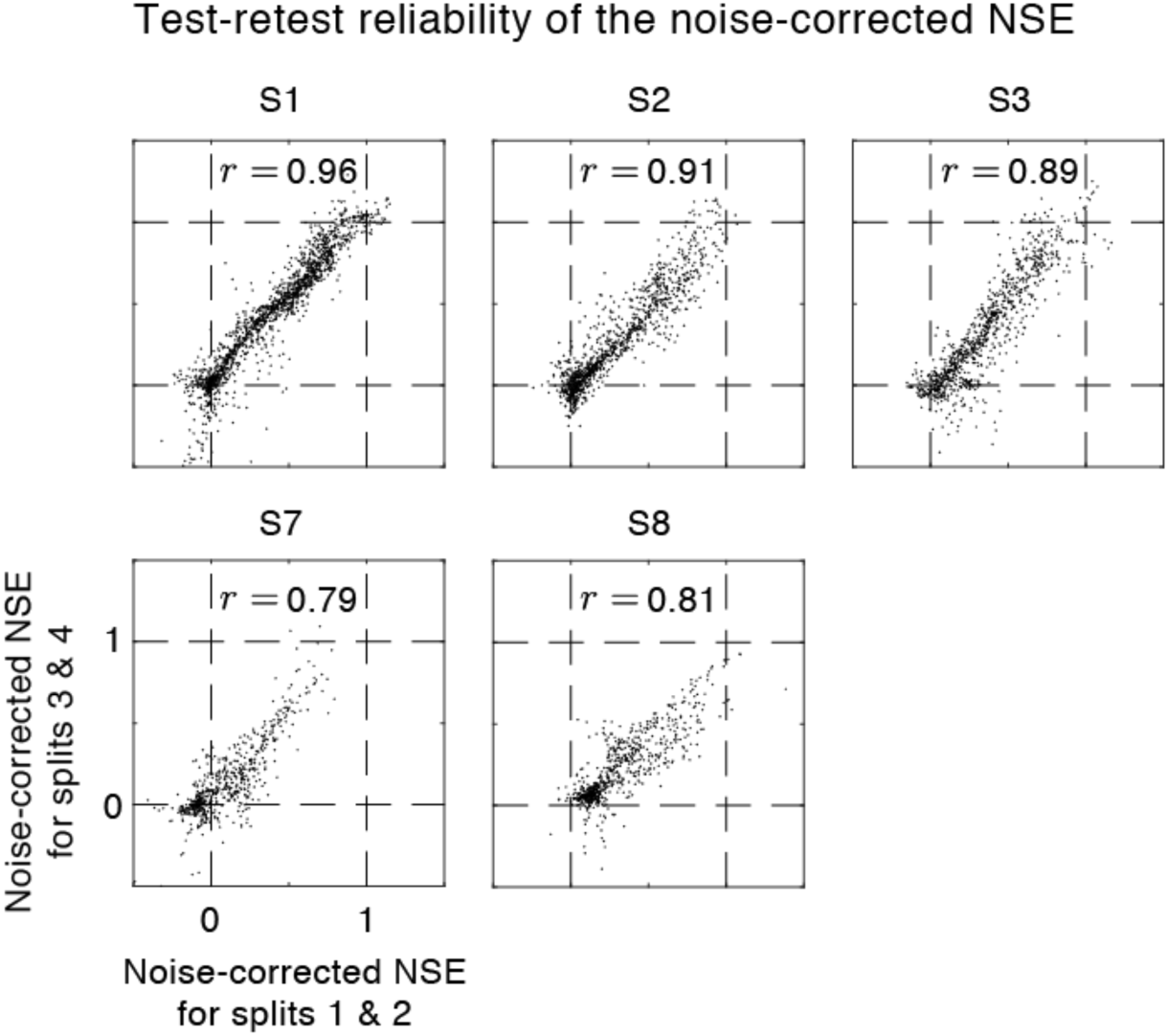
Test-retest reliability of the noise-corrected NSE (measured across splits of fMRI data). Each dot plots the noise-corrected NSE for a single voxel. The Spearman rank correlation is shown at the top of each plot for each subject. Results are shown for subjects scanned in Paradigm II, for which there was sufficient data to compute two separate estimates of the noise-corrected NSE.

**Table S1.** The presence/absence of HG duplications for each hemisphere of each subject that was scanned multiple times in the experiment. Categories were determined by inspection using the scheme described in Da Costa et al. (2011)^52^

| Subject | Hemisphere | Duplications along HG |
| --- | --- | --- |
| S1 | Left | Partial Duplication |
| S1 | Right | Single Gyrus |
| S2 | Left | Single Gyrus |
| S2 | Right | Single Gyrus |
| S3 | Left | Partial Duplication |
| S3 | Right | Single Gyrus |
| S4 | Left | Partial Duplication |
| S4 | Right | Single Gyrus |
| S5 | Left | Partial Duplication |
| S5 | Right | Partial Duplication |
| S6 | Left | Partial Duplication |
| S6 | Right | Complete Duplication |
| S7 | Left | Single Gyrus |
| S7 | Right | Complete Duplication |
| S8 | Left | Partial Gyrus |
| S8 | Right | Complete Duplication |

